# Structural Eigenmodes of the Brain to Improve the Source Localisation of EEG: Application to Epileptiform Activity

**DOI:** 10.1101/2025.07.27.667083

**Authors:** Pok Him Siu, Philippa J. Karoly, Sina Mansour L., Artemio Soto-Broceda, Levin Kuhlmann, Mark J. Cook, David B. Grayden

## Abstract

A fundamental view of neuroscience is that, in addition to neuronal activity, the structure of the brain constrains and explains brain function. An alluring formalism in computational neuroscience has been the generation of structural eigenmodes of neural activity from a matrix representing the anatomy of the brain. Traditionally, brain connectomics has been the gold standard for the coupling between structure and function. However, it has recently been suggested that simpler brain geometry can provide more explanatory power in fMRI. An adjacent modality is the source localisation problem of EEG, which aims to identify the underlying generators of EEG recordings. The underdetermined nature of the problem requires sufficient constraints to produce realistic and unique solutions of source activity. In this work, we presented a simple framework for incorporating different forms of structural brain eigenmodes to constrain the source localisation problem in epilepsy. We found that geometric eigenmodes were able to reconstruct the spread of a seizure through the brain slightly better than connectome eigenmodes, and both types of structural modes significantly outperformed commonly used approaches.

## 1 Introduction

Epilepsy is estimated to affect around 0.6% of the global population [1]. Of these estimated 50 million cases of epilepsy, around 30% are antiepileptic drug-resistant [1]. In 50% of the latter cases, the patient is diagnosed with focal epilepsy [2], where, for a carefully selected subset of patients, the treatment is to resect the culpable region of the brain, known as the epileptogenic zone. Accurate identification of the epileptogenic zone is pivotal to the diagnosis, treatment, and outcomes for these patients. The current state of the art involves patients undergoing multiday invasive studies to determine the epileptogenic zone [2]. However, methods such as stereo-electroencephalography (sEEG) are limited in their spatial coverage, come with significant risks and discomfort, and require a clear pre-implantation hypothesis [3].

Electroencephalography (EEG) and magnetoencephalography (MEG) are foundational tools in neuroscience for analysing the state of the brain non-invasively. The key advantage of these macroscopic, electrophysiological recordings are their high temporal resolution, but the trade-off is the relatively low spatial resolution of their direct measurements. A core property of EEG/MEG is that these signals are a weighted sum of neural activity and, by themselves, cannot directly define the underlying sources of neural activity.

Source imaging is a mathematical technique that aims to recreate the activity of the underlying neural sources from EEG/MEG data. However, this transformation typically involves a lower number of electrodes (∼10^2^) being mapped to a notably larger number of neural sources (*∼*10^3^-10^4^) and, therefore, is an underdetermined mathematical problem. The underdetermined nature of the source imaging problem requires sufficient constraints to produce realistic and unique solutions of source activity [4, 5]. Source imaging algorithms are hence reliant on two core components: i) a

realistic volume conduction model that solves Maxwell’s equations to convert from source space to sensor space (the ‘forward problem’) [6] and ii) mathematical constraints to provide a unique solution to the inverse transformation from sensor space to source space. Current algorithms employ biological realism with the forward model. However, the inversion constraints are typically less motivated by physiology and more by mathematical and statistical arguments [5, 7]. For instance, the minimum-norm constraint provides the lowest-energy solution, which is a plausible but not necessarily accurate approximation of neural behaviour. Constraints on the smoothness and focality of sources are also common and, although both can be said to be loosely motivated by neurophysiology, both also have a tendency to deliver biologically implausible results in practice [8].

A recent study proposed that the geometry of the brain offers a fundamental but simplistic constraint on dynamic neural activity [9]. The construction of geometric eigenmodes of the brain’s physical shape was found to offer a biologically plausible explanation of associated excitation between brain regions. Since that seminal paper, geometric eigenmodes have found varied applications in neuroscience, such as analysing the psychological effects of meditation [10], quantifying hemispheric asymmetry [11], and fingerprinting the brain to assess adolescent mental health [12]. Just as the Fourier transform decomposes complex temporal signals into a set of orthogonal sine and cosine functions, geometric eigenmodes of the brain form a spatial basis that can be used to represent distributed patterns of neural activity. Each eigenmode reflects a standing wave pattern constrained by the geometry of the brain, enabling efficient reconstruction and analysis of large-scale brain dynamics. Having such a clean modal decomposition enables geometric eigenmodes to offer a new means to constrain the source localisation problem of EEG and MEG based on our current understanding of the brain’s structure and function.

The geometry of the brain is not the only sensible structure to create these structural eigenmodes. Connectome eigenmodes, based on a matrix representation of the anatomical connectivity of the brain, may offer even more utility given that these account for both short-range local connections, as well as long-range white matter tractography. Early studies suggested that the explanatory power of geometric eigenmodes exceeded structural connectomes for explaining spontaneous and task-evoked conditions in fMRI data [9]. However, several subsequent studies have provided conflicting evidence that connectome eigenmodes are similar or superior to geometric eigenmodes [13, 14]. Given the utility of structural eigenmodes to analyse neuroimaging data, incorporating either a geometric or connectome eigenmode decomposition could improve the tractability of the inversion step in source imaging algorithms, but it is unclear which would be superior.

To understand the distinction between geometric and connectome approaches, validating novel source imaging algorithms requires a reliable source of truth; i.e., a dataset where both EEG/MEG are recorded alongside known neural sources. In practice, this ground truth is rarely available, and source imaging studies typically rely on simulated datasets or studies where the true source can be assumed from behavioural manifestations (such as the onset zone of epileptic seizures or evoked responses to stimuli). There are now two compelling reasons for creating a realistic simulation as a test data set. First, a sufficiently complex simulation enables the differentiation between geometric and connectome eigenmodes in terms of their expressiveness. Second, for the problem of source localisation in epilepsy, the underlying sources within the epileptic zone cannot be directly experimentally verified. Currently, the best measure of success is a binary classification of whether resective surgery was effective in stopping or reducing seizures [15, 16]. By introducing a realistic simulation of the underlying sources of epileptic activity with its corresponding EEG activity, a test set with a known ground truth can be used to understand the differences between different source localisation methodologies and conditions. Previous studies have typically focused on one to a few sources of various spatial extents[17, 18, 19, 20, 21, 22], but such sources are biologically questionable given that the brain is known to be a highly interconnected and complex hierarchical structure [23, 24, 25] that is capable of producing rhythmic and wavelike activity with wavelengths spanning several centimetres [9, 26, 27]. Building on successes in computational neuroscience [28, 29, 30, 31], we constructed dynamical neural mass models to produce realistic simulations of seizures.

In this study, we evaluated the utility of geometric and connectome eigenmodes as basis functions for constraining the problem of EEG/MEG source localisation. In doing so, we aimed to explore the following research questions:

1. Do geometric and connectome eigenmodes differ in their capacity to reflect neurologically meaningful activity?
2. Can incorporating such structural modes into inverse models enhance the efficiency and accuracy of EEG/MEG source localisation?
3. Does structural eigenmode source localisation show potential for improving the localisation of epileptogenic zones in patients with epilepsy?

## 2 Methods

The Methods section first describes the creation of geometric and connectome eigenmodes using real-world structural brain data, and then explains how structural eigenmodes are integrated into a source localisation method. The following sections then outline the validation datasets and evaluation metrics deployed to test the eigenmode-constrained source imaging approach.

### 2.1 Structural Brain Data

Structural connectivity data from the Human Connectome Project (HCP) [32] was used, with HCP Subject 100206 (HCP100206) as the primary subject used in this work. The T1 MRI image with skull included was used to generate surface meshes in the subject’s native space for the skin, skull, and cortex using Freesurfer’s recon-all [33]. The cortical mesh from fsaverage template space was also used to test the effects of using group-averaged structural data in the source imaging pipeline. The individual subject’s cortical mesh was aligned to the fsaverage template mesh, which consisted of 163,842 vertices per hemisphere.

Each vertex was attributed to one of the 1000 cortical regions of the Yan-1000 homotopic cortical parcellation atlas [34]. The location of each cortical region was defined as the centroid of its vertex coordinates. In addition to the 1000 cortical regions, 19 subcortical structures, defined by the cifti grayordinate template [35], were also included, resulting in a 1019-region parcellation of the entire brain.

The individual’s structural connectome was computed using a diffusion MRI tractography pipeline detailed elsewhere [36, 37] using MRtrix3 software [38]. In summary, fibre orientation distributions (FODs) were computed from tissue-specific response functions via Multi-Shell Multi-Tissue Constrained Spherical Deconvolution [39, 40]. A total of 5 million tractography streamlines were estimated by an anatomically constrained probabilistic tractography algorithm based on second-order integration over FODs [41, 42]. Notably, streamlines were randomly seeded from the gray matter-white matter interface. A radial search with a maximum radius threshold of 4 mm was used to map streamlines to the regional parcellation, resulting in a 1019×1019 connectivity matrix of streamline counts. The matrix of streamline counts was then normalised by the maximum streamline count to end in a normalised connectivity matrix with entries between 0 and 1. In addition, a 1019×1019 streamline length matrix was computed to quantify average streamline lengths between all pairs of connected regions. To obtain the group-averaged connectivity matrix of streamline counts, the sum of streamlines between regions was taken for each patient, and the aggregate matrix normalised between 0 and 1.

### 2.2 Eigenmode Creation

#### Geometric Eigenmodes

Geometric eigenmodes were constructed by solving the eigenvalue problem with the Laplace-Beltrami operator on the surface mesh of the brain [9],

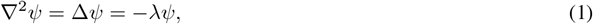

where ∇ is the gradient operator, Δ is the Laplace-Beltrami operator, and *ψ* represents the set of geometric eigenmodes with their corresponding eigenvalues *λ*. These eigenvalues and eigenmodes were ordered according to increasing spatial frequency such that *ψ*_1_ was the eigenmode with the longest wavelength. Geometric eigenmodes were also grouped into distinct eigengroups of similar wavelengths of size 2*ℓ* + 1, where *ℓ* is the *ℓ*-th eigengroup. The geometric eigenmodes were originally constructed on the 10 242 vertices white-matter surface mesh per hemisphere (icosahedron 5 spacing), and then, reduced to a 1000 region source space by averaging all vertices within each brain region of the Yan-1000 brain region parcellation [34] to match the spatial resolution of the connectome. The full set of geometric eigenmodes for the whole cortex was then the concatenation of the eigenmodes constructed for each hemisphere.

#### Connectome Eigenmodes

Connectome eigenmodes were constructed by solving the eigenvalue problem on the graph Laplacian representation of the connectome [43],

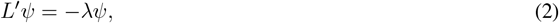

where *L*^*′*^ is the normalised graph Laplacian, a discrete version of the Laplace-Beltrami operator. This normalised graph Laplacian relates to the unnormalised graph Laplacian, *L*, through

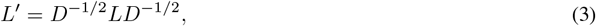

And

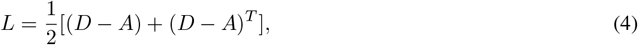

where *D* is the diagonal degree matrix and *A* is the adjacency matrix, i.e., the connectome. As with the geometric eigenmodes, the connectome eigenvalues and eigenmodes were ordered by decreasing wavelength. The 1019 brain region connectome (1000 cortical regions with 19 subcortical regions) was used as the adjacency matrix, with the 19 subcortical regions removed to obtain a 1000 region source space (comparable to the geometric source space).

### 2.3 Source Localisation

In their original form, the eigenmodes represent neural activity in neural source space. The transformation from source space to EEG measurement space can be described by a linear forward model,

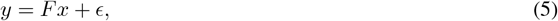

where the transformation matrix *F* ∈ ℝ^*M ×N*^ (known as the forward matrix or lead field matrix) maps any neural activity in source space *x* ∈ ℝ^*N ×T*^ to EEG/MEG sensor space *y* ∈ ℝ^*N ×T*^. The dimensions represent the number of sensors (*M*), the number of possible sources in the brain (*N*), and the number of time steps (*T*), while *ϵ* is a matrix representing the overall measurement noise for each time step.

For this study, *F* was constructed with a head volume conduction model computed using a three-shell Boundary Element Method (BEM) [44] approach implemented in OpenMEEG and MNE-Python [45, 46]. *M*, the number of sensors, varies dependent on the EEG-spacing montage used, such that 10-20 spacing involves 21 electrodes, 10-10 spacing involves 88 electrodes, 10-05 spacing involves 339 electrodes. Unless specified, the 10-10 spacing montage was used. *N*, the number of sources, was set to 1000 dipoles centred at each cortical region in the Yan-Atlas and representing an average spacing close to 1 cm.

Applying the lead field matrix, *F*, to the set of structural eigenmodes, *ψ* ∈ ℝ^*N ×K*^, provided a corresponding set of *K×*EEG modes. Hence, Eq. 5 becomes

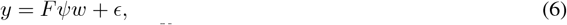

where *Fψ* represents the matrix of EEG modes (*M × K*) and *w* ∈ ℝ^*K*^ is the corresponding weight vector. The least squares solution to Eq. 6 is

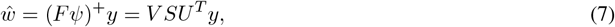

where the + operator denotes the Moore-Penrose pseudoinverse, obtained via the singular value decomposition (SVD) into rotational matrices *U, V* ^*T*^ and a diagonal non-negative real matrix *S*. It is known that lower spatial modes dominate cortical activity [9], so we biased the eigenmodes 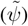 by normalising by the Froebius norm and then multiplying with a diagonal weighting matrix (*W*), whose elements are defined by

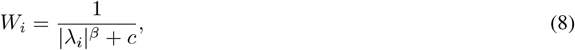

where *c* is a constant term to ensure numerical stability and |*λ*|^*β*^ is the mean of the absolute value of the *i*-th eigenvalue to the power of *β*. For the geometric eigenmodes, we set 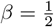 such that the weighting was 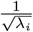. The rationale lies from the spherical harmonic equations, where 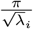 represents the normalised spatial wavelength of the *i*-th eigenmode expressed as a fraction of the head diameter. Hence, this weighting aligns with the low-pass filtering properties of EEG through the physical meaning of the normalised spatial wavelength and can be seen as a proxy for the fraction of EEG sensors that can meaningfully capture each eigenmode (see Fig 3). The constant *c* can then be set naturally as 2*π*, since the geometric eigenmodes are defined on a hemisphere, resulting in the weight of the 0^th^ (constant) eigenmodes being 1*/*2, representing half of the head. For the connectome eigenmodes, the eigenvalues do not have an explicit spatial interpretation in metric units, so we set *β* = 1 and 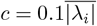, where 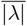 is the mean of the absolute value of all eigenvalues, resulting in weights that closely align with the geometric eigenmode weights after normalisation (see Fig S3).

**Figure 1.**
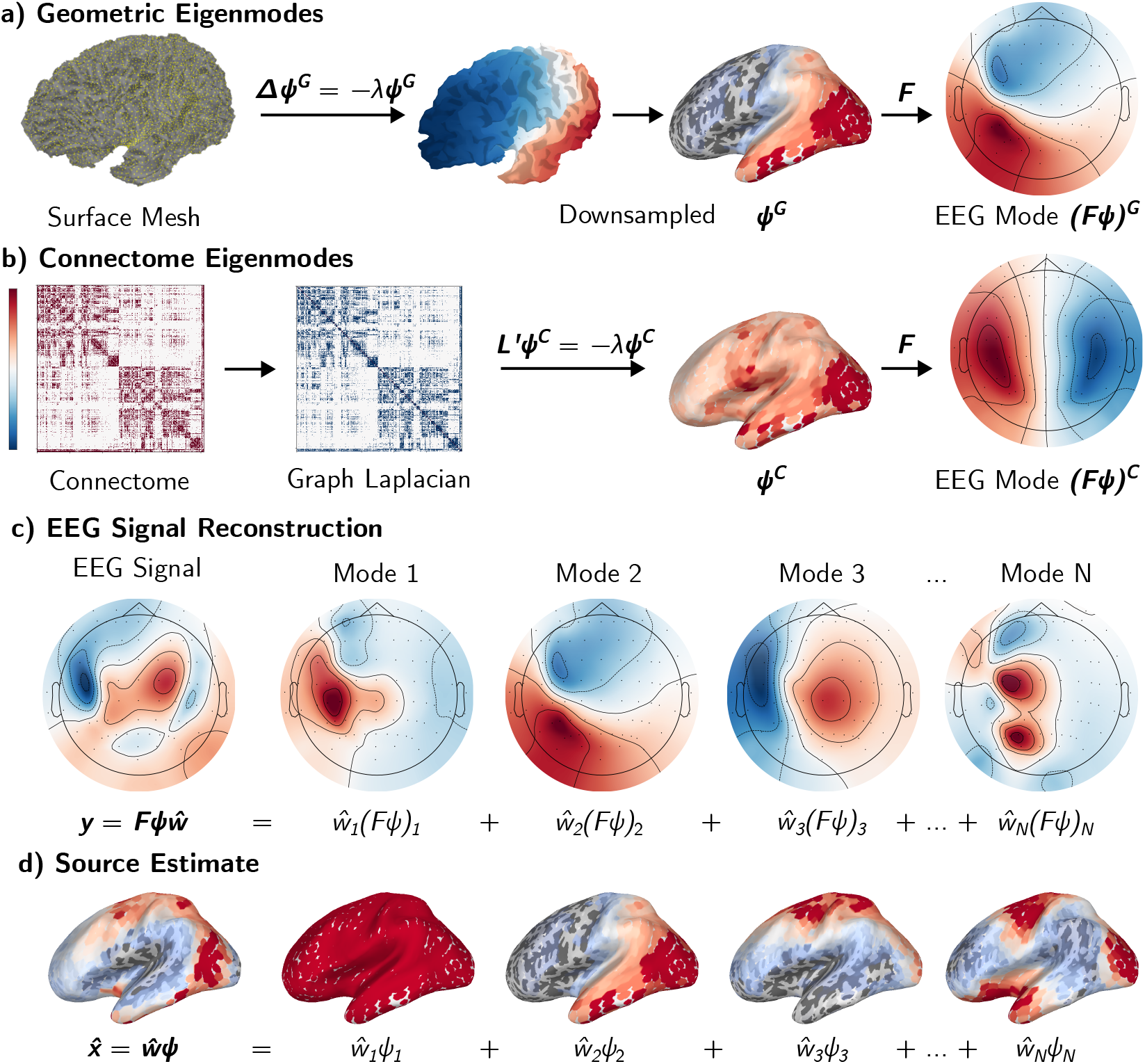
Creation of Structural Eigenmodes and Source Estimation. a) Following Eq.1, geometric eigenmodes (*ψ*^*G*^) are generated by solving the eigenvalue problem of the Laplace-Beltrami operator on the surface MRI mesh. This is downsampled to a 1000 discrete cortical region space. The forward operator (*F*) is applied to transform the eigenmode from source space to EEG space to produce a corresponding set of EEG modes. b) Following Eq.2, connectome eigenmodes (*ψ*^*C*^) are directly generated in the 1000 cortical region space by solving the eigenvalue problem on the graph Laplacian representation of the 1019 region connectome and then truncating to cortical regions only. c) Following Eq.6, the EEG signal can be decomposed into a weighted sum of the different EEG modes. The aim is to find the weights 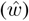 as given by Eq.7. d) Finally, the source estimate 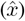 can be constructed as a weighted sum of the estimated weights and the corresponding structural eigenmodes in source space.

**Figure 2.**
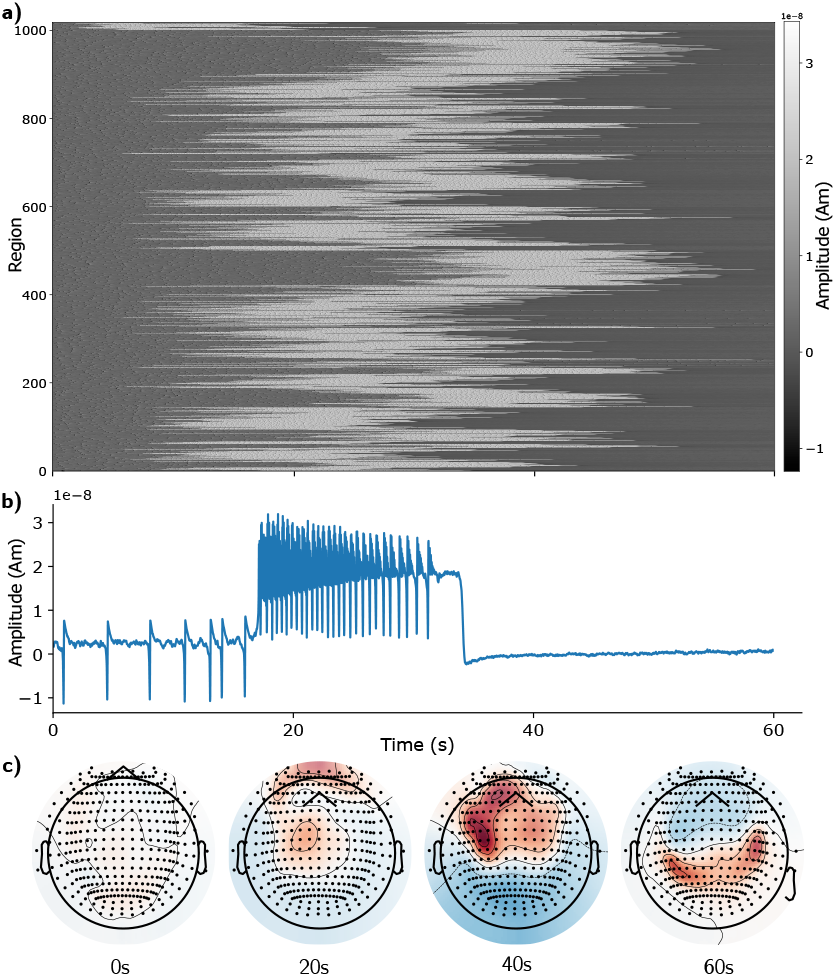
Example Seizure Simulation of the Epileptor Model. a) Carpet plot [29] of a seizure simulation starting from the thalamus with connectivity strength, *g* = 2. The areas in white signify that the region is undergoing high-energy, epileptic activity. At *g* = 2, the seizure spreads throughout the entire brain. b) Example time series of the Epileptor model. Key features are the interictal spikes (which go to ∼ −10 nAm), the spike-wave discharges at ∼20 nAm, and the lack of any interictal spikes in the refractory period afterwards. Please refer to the original paper for a comprehensive comparison of the simulated time series with invasive recordings in zebrafish and human [48]. Note that, as per Eq.11, the original equations were adjusted such that the time series have baseline at around 0 and the seizure state is positive. c) Example of EEG topography. Using the forward model (Eq.5), the source-level simulations from panel a) were projected to the sensor space to generate EEG topographies. The corresponding EEG trace is shown in Fig S2. A neurologist reviewer confirmed that the simulated EEG exhibited clinically consistent features, including focal onset, spatial spread, and temporal evolution with characteristic slowing.

**Figure 3.**
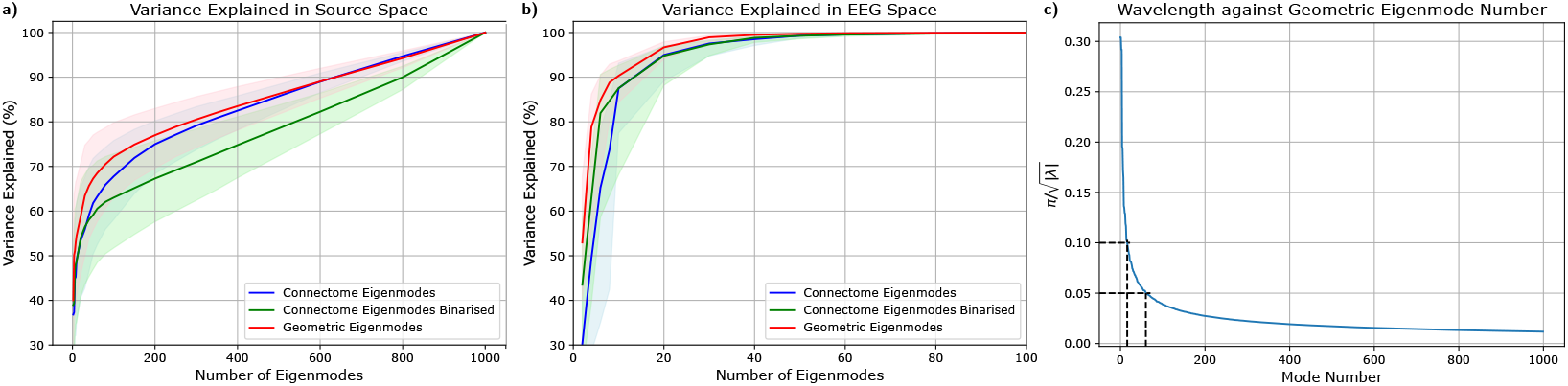
Reconstruction of activity at source and sensor level. For subfigures a) and b), variance explained is graphed against an increasing number of eigenmodes used in the reconstruction. No weighting was applied to the eigenmodes. The blue line indicates connectome eigenmodes, green indicates connectome eigenmodes generated with a binarised version of the connectome, and red indicates geometric eigenmodes. The shaded regions indicate the range of variance explained across all simulations. a) Variance explained in source space shows how well each structural eigenmode type is able to reconstruct the source simulation from the Epileptor model. b) Variance explained in EEG space indicates how well each structural eigenmode type is able to reconstruct the simulation after it is converted into EEG space using a forward model. The EEG montage used was 10-05 spacing (343 electrodes) for explanatory purposes. Note the difference in x-axis limits due to how quickly the variance explained reaches 100% in EEG space. c) The wavelength of the geometric eigenmode represented as a proportion of the head dimension. The 0^th^ eigenvalue is ignored. The dashed lines correspond to common EEG montages of 10% and 5% spacing. The corresponding mode numbers are 16 and 60.

Finally, 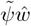 represents the weighted sums of the biased structural eigenmodes, so everything was combined to yield the aggregate estimated source activity,

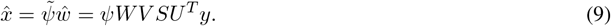

It is important to consider the singular values of the EEG modes due to the effects of noise and modal amplification [47]. Small singular values apply a large weight to their respective component, leading to any perturbations being significantly amplified. Under the SVD framework, small singular values in *S* are used to determine modes to truncate in *U, V*, referred to as the ‘rcond’ parameter in numpy.linalg. Truncation performs a similar role to the regularisation parameter (*α* or *λ*^2^) in traditional regularisation approaches to deal with noise. Unless stated, we set the threshold to 10^−*SNR/*10^, based on the typical regularisation value used in minimum-norm approaches.

### 2.4 Simulated Evaluation Data

Simulations of seizure activity, generated from connected neural mass models, were used to represent neural activity at the source level. Using the previously mentioned forward model (see Eq. 5), the “ground truth” neural source activity was converted into EEG/MEG space. As a result, a simulated dataset, consisting of both macroscale electrophysiological data as well as a corresponding ground truth, could be used to evaluate our source localisation approach.

The interconnected neural mass models were defined according to the Epileptor equations [48, 49], an established model that has been widely used to describe the source and spread of epileptic activity with direct usage in clinical trials (EPINOV) [31, 50]. The simulation pipeline is described below.

#### Epileptor Equations

The most common implementation of the Epileptor model, as implemented in the Virtual Brain Project (TVB) [51], characterises the dynamical behaviour of epileptic seizures using five state variables (*x*_1_, *y*_1_, *x*_2_, *y*_2_, *z*) plus a dummy variable (*u*) within region *i* [49]. The equations are:

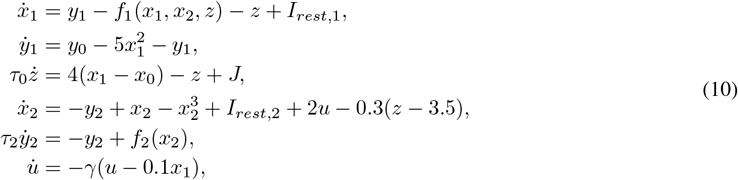

where:

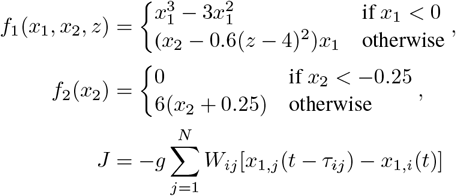

and *x*_1_(*t*), *y*_1_(*t*) govern the rapid discharges on the fast timescale, *x*_2_(*t*), *y*_2_(*t*) govern spike-and-waves on the intermediate timescale, *z*(*t*) is the permittivity variable that operates on a slow timescale affecting the transition between seizure and non-seizure states, *u*(*t*) is a dummy variable for low-pass filtering signals from *x*_1_ to *x*_2_, *x*_0_, *y*_0_ are threshold constants, *τ*_0_ and *τ*_2_ are time constants, *I*_*rest*,1_ and *I*_*rest*,2_ are constant injection currents, *γ* is the time constant of the low-pass filter, *J* is the current coupling into region *i* from other regions *j, g* is a global coupling strength parameter, and *W*_*ij*_ are the coupling weights as defined in the structural connectivity matrix into region *i* from region *j*.

These equations are used to generate time series that represent neural activity at each brain region. To represent extracellular dipole sources (*x*), the above equations are modified to become

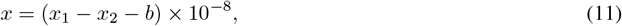

where *x*_1_ − *x*_2_ is used instead of *x*_2_− *x*_1_, as in Jirsa et al. [48], to ensure that the seizure state has a positive orientation, and 10^−8^ converts to units of Am. Notably, *b* is a baseline constant, subtracted from the Epileptor model in its original form, to ensure dipole sources maintain approximately zero baseline activity, as expected from the Debye shielding of ions in the electrolyte solution of the extracellular space [4].

#### Seizure Simulation

The simulations used 1019 coupled masses of the Epileptor models, as defined in Equation 10, through the coupling current *J*. The connectivity weight matrix *W*_*ij*_ was defined by the 1019 region connectome (see Sec. 2.1) on a specific subject (HCP100206). A Heun’s stochastic integrator was used with Gaussian additive noise (∼*N* (0, *σ*^2^)) added to the variables *x*_2_ and *y*_2_ [30, 48]. An integration step size (Δ*t*) of 0.05 was used. To match the temporal properties of focal seizures, the timescale of the model output was scaled such that 256 time steps correspond to 1 s as similarly done by Jirsa et al. [49]. Seizure simulations were chosen to last 1 minute to allow sufficient epileptogenic activity to develop.

To generate each simulation, a single given region was first defined as the ‘epileptogenic zone’. All regions other than the epileptogenic zone were set to have excitability of *x*_0_ = −2.1, i.e., tuned to be at the cusp of a bifurcation [30, 52]. To trigger a seizure, the excitability (*x*_0_) of the specified epileptogenic zone was increased to −1.6. The global coupling strength (*g*) was set to *g* = 1.5 or *g* = 2 to obtain two different degrees of seizure spread. In total, 2038 simulations were generated to produce a dataset with variations in seizure morphology by changing the specified epileptogenic zone and the global coupling strength.

#### Mapping to Sensor Space with MNE Python

The source space was defined on the cortical surface such that the centre of each cortical region of the Yan-1000 parcellation was approximated as a dipole oriented normal to the cortical surface. Seizure (Epileptor neural mass model) simulations were mapped directly to source space based on the cortical parcellation of the subject (either HCP100206 or fsaverage). These source space simulations were then mapped into EEG sensor space using OpenMEEG’s [45] implementation of the Boundary Element Method (BEM) for the forward problem with the corresponding forward model for the subject. The BEM model consisted of three concentric layers, with standard conductivity values assigned as 0.3 S*/*m (skin), 0.006 S*/*m (skull), and 0.3 S*/*m (cortex) [46]. For solving the inverse problem, the group-average forward model may be used to replicate a real-world scenario where the forward model is not known. The number of sensors varies depending on the EEG-spacing montage used, such that 10-20 spacing involves 21 electrodes, 10-10 spacing involves 88 electrodes, 10-05 spacing involves 339 electrodes. The current study derived all the results using EEG; however, the described analyses and code perform similarly for MEG.

### 2.5 Evaluation

#### 2.5.1 Simulation Metrics

Four evaluation metrics were used to assess the accuracy of source localisation within the simulations. These metrics are as follows:

##### Cosine Score

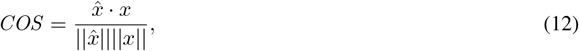

where *x* is the true signal and 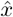 is the source estimate. Depending on the figure, this may be presented for each time point or averaged across time. Cosine score captures relative distributional accuracy, reflecting how well the relative pattern of activation across the source space matches the true activity.

##### Normalised Residual Sum of Squares

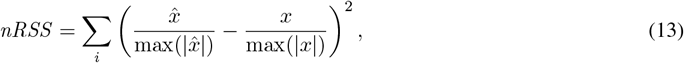

where the data was normalised with the max to compare across methods with differing units.

##### Variance Explained

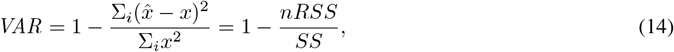

where *nRSS* are the residual sum of squares and *SS* are the sum of squares over all possible sources *i* in source space, and the max normalisation is performed as above. We note that, for our specific signal, a baseline constant was already identified and subtracted (see Eq.11) such that the baseline for *x* was at 0.

##### Region Localisation Error

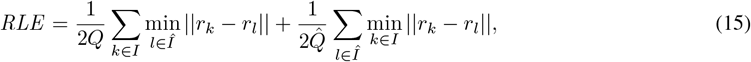

where *I* and *Î* represent the true and estimated indexes of active sources, respectively, *Q* and 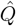 represent the number of true and estimated active sources, and *r*_*k*_ denotes the position of the *k*-th dipole source in space as per MNE-Python’s implementation [46]. Sufficiently active sources are algorithmically identified to be beyond Otsu’s Threshold [53], as suggested by Sun et al. [15, 28]. The region localisation error was computed across time as well as at 50% of maximum EEG power, representing a sensible extent of seizure spread and the rising phase of a seizure [16]. Region localisation error captures spatial accuracy, reflecting how precisely the estimated source activity is positioned in space.

#### 2.5.2 Simulation Test Conditions

To ensure experimental rigour, we performed four complementary evaluations, designed to test the comparative performance, biological plausibility, and robustness of the structural eigenmodes approach. Statistical significance was assessed with a paired Wilcoxon signed-rank test between the geometric eigenmode approach and alternative approaches. These test conditions are as follows.

1. Benchmarking with other algorithms To benchmark, we compared different off-the-shelf source localisation algorithms (dSPM, MNE, sLORETA and eLORETA) on our dataset as implemented in MNE-Python [46].
2. Surrogate mode testing We applied a surrogate mode test to assess whether the original eigenmodes performed significantly better than structurally similar alternatives. Surrogate modes were generated using two approaches: roll, which applies a cyclic shift to all eigenmodes using NumPy’s roll function (preserving orthogonality), and rotation, which applies orthogonal rotation matrices within each harmonic group, preserving their internal spatial structure, following the method of Koussis et al. [54]. For each simulation, we computed a performance score for the original eigenmodes and compared it to the distribution of surrogate scores. The percentile rank of the original mode relative to the surrogate distribution was calculated for each simulation. To assess whether the original modes consistently outperformed surrogates across simulations, we applied a paired Wilcoxon signed-rank test on the percentile values (centred around chance level by subtracting 50).
3. Subject-specific vs. group-average comparison The inverse crime method [55] is referred to as such when the same model is used for both the forward and inverse solutions. To test the extent of this, we tested simulations where the signal in Eq.5 was generated with the subject-specific forward model and the inverse used a group-average forward model (fsaverage); i.e., the EEG modes in Eq.6 were created using the group-average forward. We also tested the effect of using fsaverage to generate the geometric eigenmodes and a HCP group-averaged connectome to generate the connectome eigenmodes.
4. Varying signal-to-noise conditions for EEG EEG is well-known to be a noisy signal in practice. To simulate the challenge of noise contamination, Gaussian white noise was added to the EEG signal with a signal-to-noise ratio of 3 dB, 10 dB, or None (signifying no white noise added). The standard deviation of the noise was given by 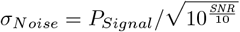.

For a more direct comparison with previous literature, which focuses typically on one to few sources of various extent [17, 18, 19, 20, 21, 22], the interested reader can refer to the evaluation on a simple set of single-patch simulations in Sec. S1.1. These patch simulations are also more analogous to the interictal spikes analysed in the following section with real patient data. In addition, to assess whether the results obtained in the lower resolution source space held at a higher spatial resolution, further testing was conducted using a source space consisting of 10 242 dipoles per hemisphere (icosahedron 5 spacing), with eigenmodes computed at this resolution (see Sec. S1.3).

#### 2.5.3 Real Data

With simulation, we tested the eigenmode approach to source localisation under different conditions and identified how best to use the approach. Next, the eigenmode approach was validated in a cohort of 20 drug-resistant, focal epilepsy patients who had undergone resective surgery to obtain seizure-freedom as per ILAE-1 or ILAE-2 classification. The data [15] had high-density 76-channel scalp EEG recordings where 5 to 90 interictal spikes were identified and averaged per patient. All data processing was performed by the data creators. Averaged interictal spikes per patient were input into the eigenmode algorithm with a group-average model and then compared to the clinical records of resected area. A signal-to-noise ratio of 10*log*_10_(*N*), where *N* was the number of interictal spikes, was assumed for each patient.

The resected region was mapped onto the group-averaged source space (fsaverage) and treated as the base truth. This ‘ground truth’ was compared to the estimate produced by the geometric/connectome eigenmode source localisation approaches using geometric/connectome eigenmodes generated from group-averaged data.

Given that the task was a binary classification, for better comparison with the original paper, we evaluated the following metrics in this section: region localisation error (RLE), precision, recall, and the area under the receiver operating characteristic curve (ROC-AUC). Let *TP* denote true positives, *FP* false positives, *FN* false negatives, and *TN* true negatives. Then, precision and recall are defined as

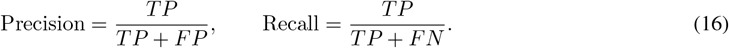

As with simulation, Otsu’s threshold was used to determine the appropriate threshold to use for these threshold-based metrics.

In contrast, the ROC-AUC evaluates the ability of the classifier to discriminate between positive and negative classes across all possible thresholds. The ROC curve plots the true positive rate 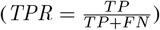 against the false positive rate 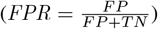, and the AUC quantifies the area under this curve. A perfect algorithm achieves an AUC of 1, while a random algorithm has an expected AUC of 0.5.

## 3 Results

### 3.1 Structural Eigenmodes Represent Simulated Cortical Source Activity

First, it is vital to confirm that simulations of ictal activity can be decomposed into structural eigenmodes. This is demonstrated by Figure 3, which shows that a relatively small number of eigenmodes were required to reconstruct the simulations in both EEG and source space.

In terms of source-space reconstruction (Fig 3a), both geometric and connectome eigenmodes were able to represent a significant portion (75% and 77%, respectively) of the simulated source signal using 200 eigenmodes. The results were consistent with previous work using structural eigenmodes to reconstruct fMRI data, which is more analogous to source-space than EEG signals, with >80% reconstruction accuracy using 200 eigenmodes for a single hemisphere [9]. It is interesting to note that, whilst it is common to binarise the connectome before using it, we find that the binarised connectome had poorer explanatory power in source-space than the non-binarised connectome eigenmodes, agreeing with recent articles that binarisation may be too simplistic and removes valuable information [56]. At low numbers of eigenmodes, geometric eigenmodes appeared to have higher explanatory power than connectome eigenmodes (at 50 eigenmodes, 62% vs. 77%). This gap then disappeared as more eigenmodes were introduced.

In terms of EEG-space reconstruction (Fig 3 b), under no-noise conditions, both connectome eigenmodes and geometric eigenmodes were able to reconstruct beyond 90% of the variance of an EEG signal with only 20 eigenmodes across both hemispheres. Performance quickly reached 100% variance explained, reflecting the high correlation between EEG electrodes and the presence of low singular values in the SVD of the EEG-mode space. This trend, where the variance explained quickly saturates, aligns with the physical interpretation of the eigenvalues of the geometric eigenmodes as spatial frequencies, as demonstrated by Fig3c). More specifically, the spatial wavelength is constructed through the relation 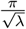, which can be interpreted as modes that span the same spacing as the EEG electrodes. A spacing of 5%, as per a 10-05 montage, would be expected to saturate in complexity at around 60 modes, as demonstrated by Figs.3b and c.

### 3.2 Structural Eigenmodes Enable Source Localisation of Seizure Spread

Next, we compared the accuracy when geometric and connectome eigenmodes were used to predict the underlying neural sources. The following sections present a detailed evaluation of the performance of the structural eigenmode approach when using a different number of modes, under different signal to noise ratios, and when compared to other approaches. Figure 4 provides a visual representation of an example simulation, where both structural eigenmode approaches were able to track the spread of a seizure originating from the thalamus.

**Figure 4.**
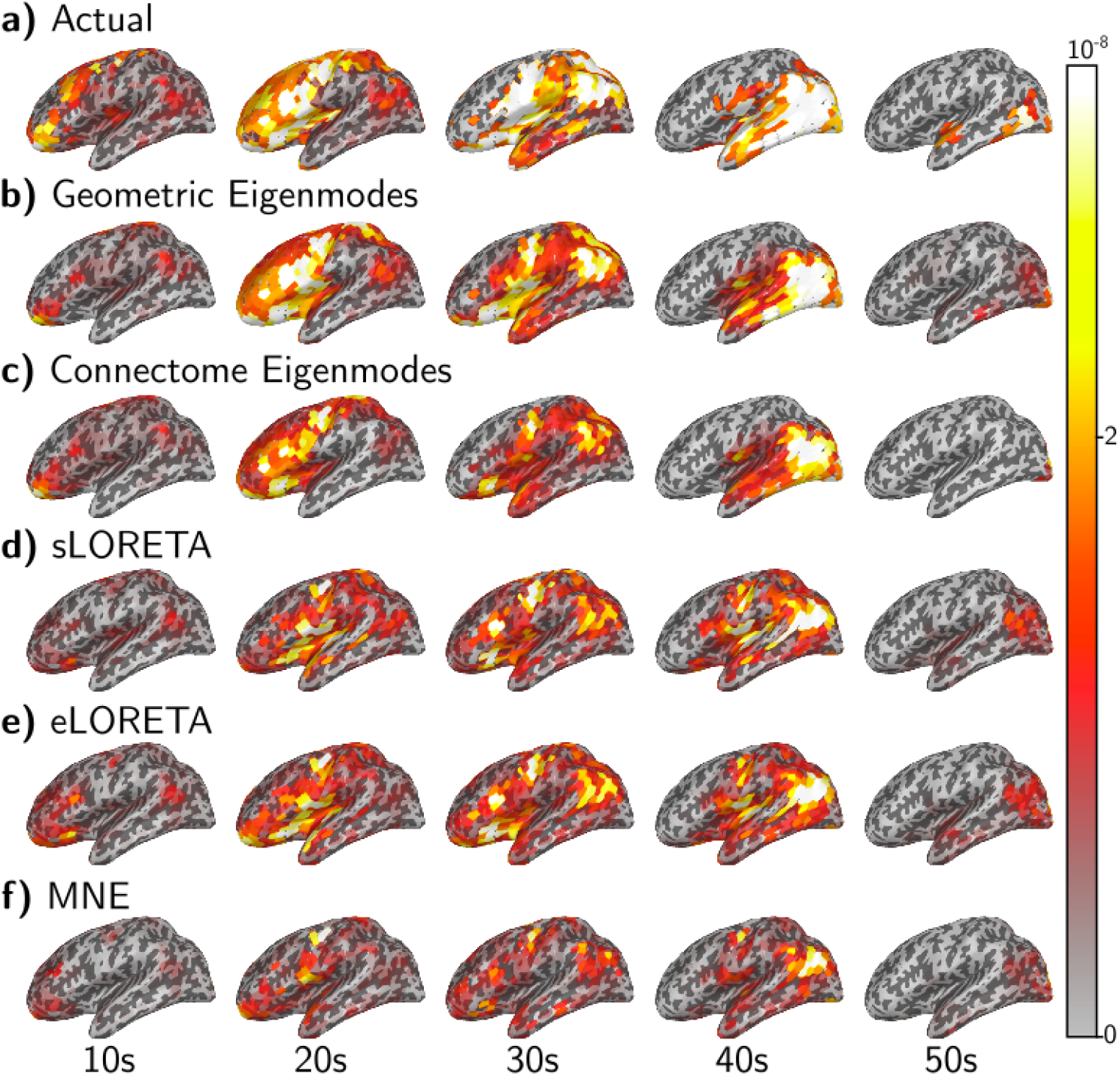
Source localisation example for seizure originating from thalamus plotted on left hemisphere. Five time points are selected in increments of 10 s and visualised on the left hemisphere. For the right hemisphere result, please see Fig S12. 50% EEG power for this simulation occurs at the 30 s. The colour scale is set such that maximum amplitude is white, progressing to yellow, orange, red, and transparent for 0. a) The actual or ‘ground truth’ source activity generated from the Epileptor model. As time progresses, the seizure spreads from the anterior to posterior regions of the brain. During this time, different areas of the brain are activated in varying spatial extents and patterns. b) & c) Both geometric eigenmodes and connectome eigenmodes are able to perform reasonably well in sparse activations (10 s and 50 s), as well as dispersed activations (20 s to 40 s). d)-f) The LORETA family predict more focal sources with regularisation parameter set according to the 10^−*SNR/*10^ heuristic. For visualisation and to improve performance, the orientation of the dipole was allowed to be loose for the LORETA family, which we demonstrate to be near equivalent to taking the absolute value of a fixed orientation estimate. As such, the performance increase for the LORETA family can be considered misleading.

#### 3.2.1 Performance against number of modes — the over-vs under-parameterised regimes

Figure 5 depicts the effects on localisation performance as the number of modes increases. With no simulated EEG noise and no truncation, increasing the number of modes initially improves most metrics (RLE at 50% power and cosine score), before reaching a peak and deteriorating. For the cosine score and for geometric eigenmodes, this local peak is at 32 modes, corresponding to 4 eigengroups per hemisphere. All metrics show progressively worse performance, until a local maxima/minima, corresponding to 88, the number of electrodes under the standard 10-10 EEG layout. After reaching 88 electrodes, performance then begins to improve until it reaches a plateau. This trend is heavily related to the ‘Double Descent’ phenomena seen in machine learning (as discussed in Sec.4) [57, 58].

**Figure 5.**
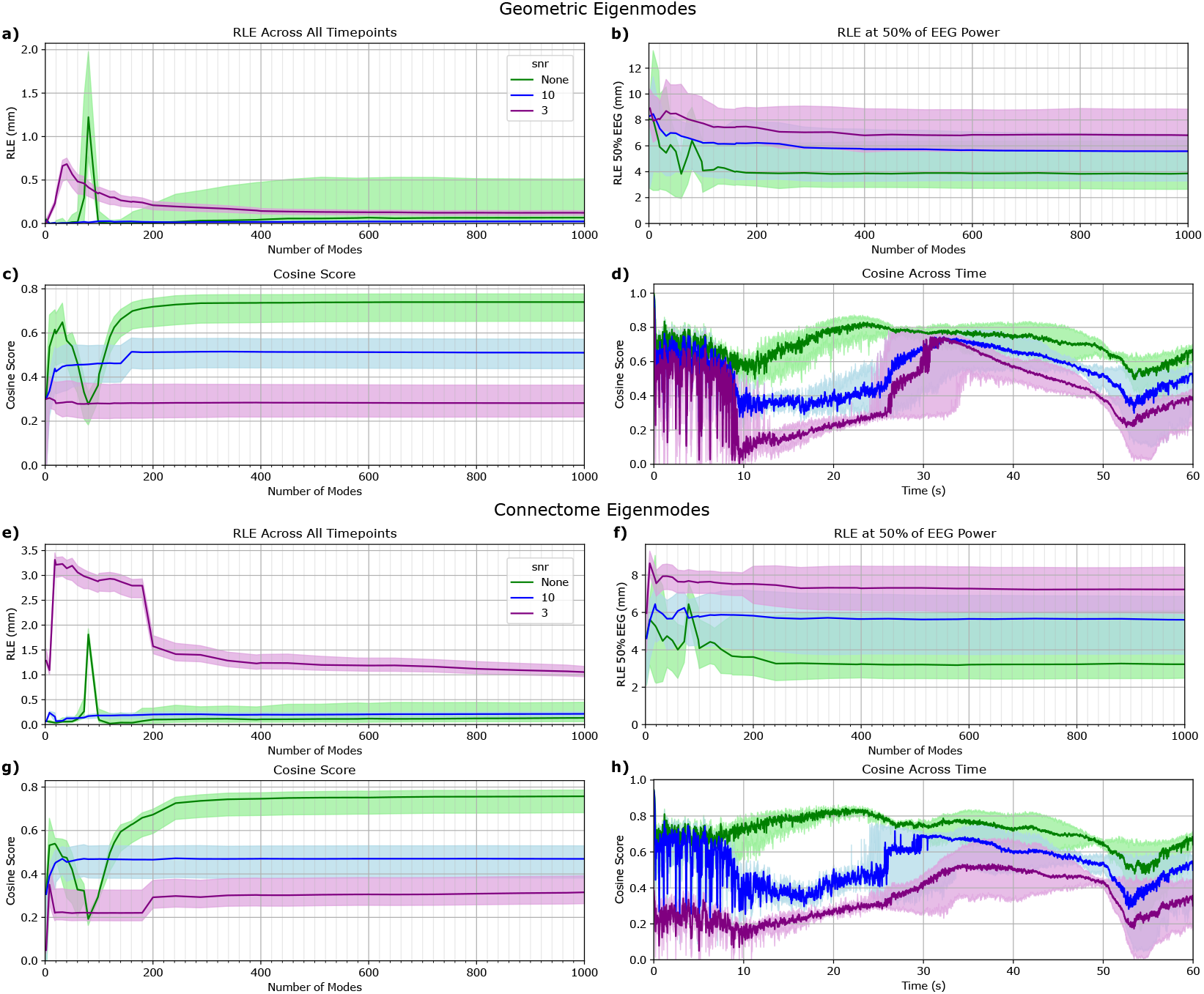
Geometric and connectome eigenmode metrics against number of modes and signal to noise conditions. Four different metrics: a) & e) region localisation error averaged across all timepoints, b) & f) region localisation error at 50% of EEG power, c) & g) cosine score, d) & h) cosine score averaged across all timepoints, are shown for both the geometric and connectome eigenmode approaches. The different coloured lines represent different levels of EEG noise added to the simulation. A SNR of ‘None’ signifies no noise was added (green), ‘10’ means a SNR of 10 dB (blue), and ‘3’ means a SNR of 3 dB (purple). The shaded region about the line signifies the interquartile range for that metric across all simulations. The solid line represents the median value of the metric across all simulations for the specific number of modes. For cosine across time, the maximum across all number of modes is taken first before the median and interquartile range is calculated for the figure. The plots shown are for 10-10 EEG spacing, consisting of 88 electrodes.

Comparing geometric and connectome eigenmodes, both types of structural eigenmodes appear to perform similarly well under all noise conditions but with some notable differences. For RLE at 50% of EEG power, connectome eigenmodes appear to be slightly better under no noise conditions by 1 mm. For averaged cosine score, geometric eigenmodes reach the optimal under a lower number of modes, mirroring expectations from Fig 3a. For cosine across time, before the 10 second mark when the seizure has not spread to too many regions, geometric eigenmodes had a higher score under high noise conditions. This is likely due to the geometric eigenmodes using a fewer number of low-wavelength modes to explain the data and thus slightly less affected by noise.

These results were consistent when using EEG 10-05 spacing consisting of 339 electrodes and EEG 10-20 spacing consisting of 21 electrodes, and when both the geometric and connectome eigenmodes were used in conjunction (see Sec S1.2&S1.4). Overall, the results of this section suggest that there are two sensible choices for the number of modes, whether it be the under-parameterised or over-parameterised regime, provided the choice is far from the number of electrodes of the EEG montage. The remainder of the presented results used 1000 eigenmodes (unless otherwise stated) to take advantage of over-parameterisation.

#### 3.2.2 Testing, validation, and comparison to other approaches

The performance of different variations of the source location approaches is shown in Figure 6, with the mean and standard deviation provided in Table S1. Under high noise conditions (*SNR* = 3 dB), all methods performed relatively poorly, with the cosine score having a large deviation and below 0.40. Therefore, the evaluation was conducted under optimal and medium (*SNR* = 10 dB) noise conditions. Statistically significant differences in performance metrics between the geometric eigenmode approach and alternative methods were evaluated using signed-rank test with a p-value *<* 0.01.

**Figure 6.**
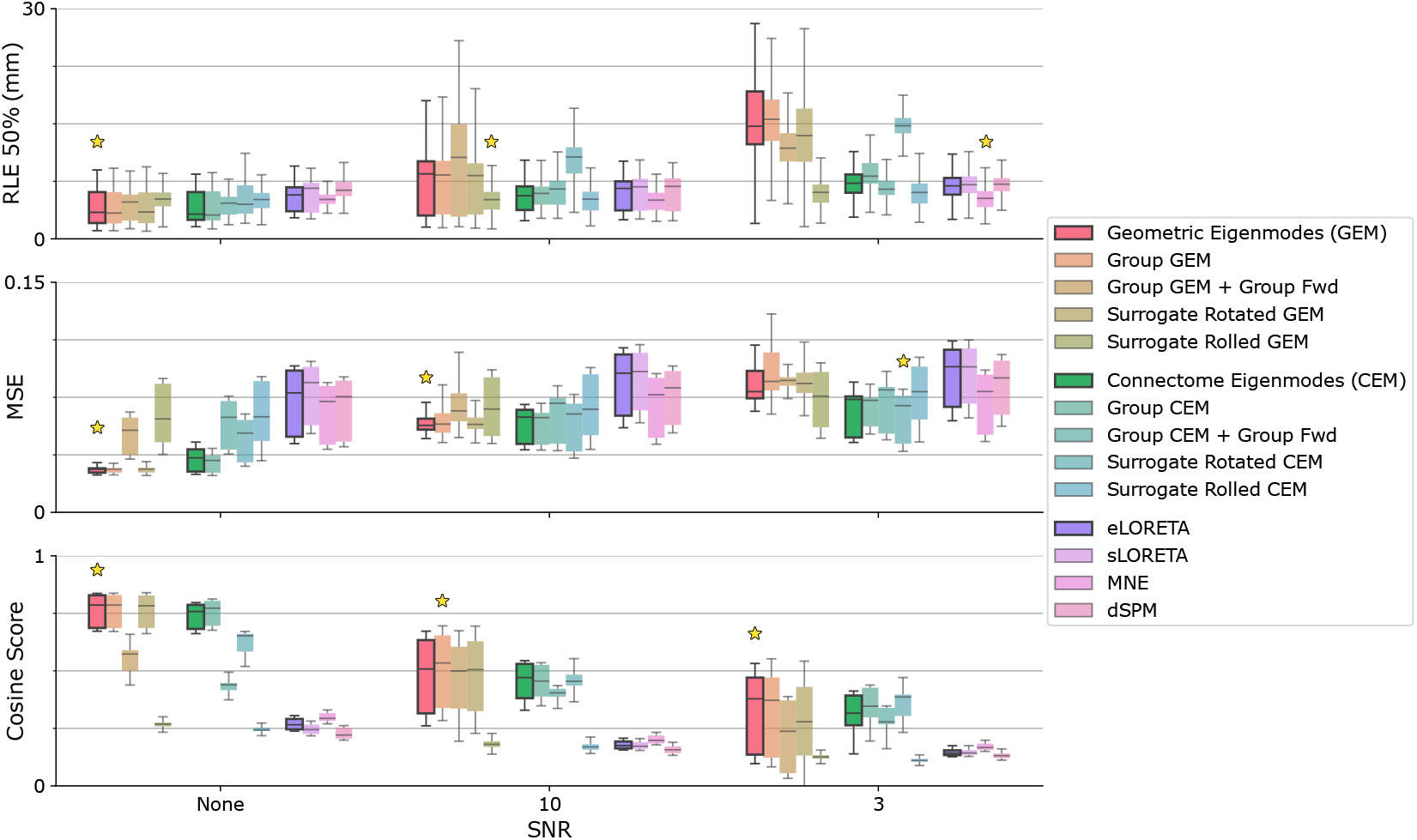
Comparisons of the structural eigenmode approach against all other methods and surrogate testing. Source localisation algorithms are grouped based on their type: GEM (geometric eigenmodes), CEM (connectome eigenmodes), and comparison methods. The highlighted boxplots correspond to the base method for geometric eigenmodes, connectome eigenmodes, and eLoreta for a standard comparison under normal conditions. Three metrics are graphed: i) region localisation error at 50% EEG power, ii) normalised mean squared error, and iii) cosine score. The star above the boxplot signifies the algorithm with the best performance for that metric and SNR. For geometric and connectome eigenmodes, eigenmodes generated using group-averaged structural data were tested. The scenario when the subject’s MRI scan was not available (where the group-averaged eigenmodes and group-averaged forward model were used together) was also tested. Surrogate eigenmodes were generated for the geometric and connectome eigenmodes as explained in Sec.2.

**Figure 7.**
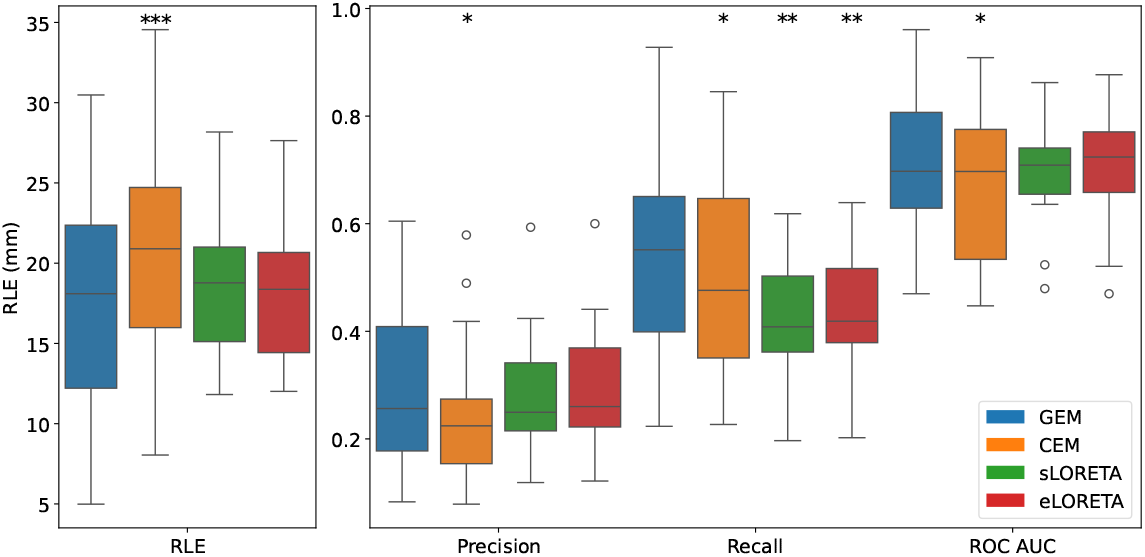
EEG 10-10 spacing clinical validation in drug-resistant epilepsy patients in comparison with surgical resection. The boxplots demonstrate the range of values for the different metrics and source localisation algorithms. The whiskers extend to the minimum and maximum values, except for algorithmically determined outliers, which are denoted as circles. The box contains the interquartile range, with the horizontal line inside the box denoting the mean. For GEM region, localisation error was 17.41*±*7.77 mm, precision was 0.29*±*0.16, recall was 0.56*±*0.17, and ROC AUC was 0.72 0.15. For CEM region, localisation error was 20.76*±*6.52 mm, precision was 0.25*±*0.14, recall was 0.50*±*0.19, and ROC*±*AUC was 0.67 0.15. The asterisks (*) above the boxplots indicate the level of statistical significance between the geometric eigenmode approach and the corresponding method, with \**p <* 0.05, \*\**p <* 0.01, \*\*\**p <* 0.001. Recall for GEM was found to be significantly higher than sLoreta and eLoreta at an *α* = 0.01. The other metrics had no statistically significant differences and were comparable.

##### Performance of established methods

For all signal-to-noise ratios, the established methods (dSPM, MNE, sLORETA, eLORETA) were unable to track the progression of a seizure in the simulated dataset. This is evidenced by the relatively poor cosine score and variance explained at around 0 or lower. The primary issue is that the methods were unable to determine the correct polarity of one-sided simulations, wherein seizure states were modelled as positive deflections from the baseline. Sec. S1.5 demonstrates that by applying a loose orientation or by taking the absolute value of the estimates, the metrics increase substantially, but this can be considered a misleading improvement given that the polarity of the estimate is known to be incorrect. Sec. S1.4 displays that using 10-05 EEG spacing (343 electrodes), the established methods had a significant increase in performance and were able to determine the correct polarity of the underlying sources.

##### Comparison between structural eigenmode approaches

Under optimal no-noise conditions, geometric eigenmodes offered statistically significant performance improvement over connectome eigenmodes. Time-averaged region localisation error was 0.12 *±* 0.14 mm vs. 0.24 *±* 0.20 mm, region localisation at 50% EEG power was 4.05 ± 2.08 mm vs.

4.20 *±* 1.85 mm, cosine score was 0.76 *±* 0.07 vs. 0.74 *±* 0.05, mean squared error was 0.028 *±* 0.002 vs. 0.034 *±* 0.007, and variance explained was 0.58 *±* 0.10 vs. 0.50 *±* 0.04 for geometric eigenmodes vs. connectome eigenmodes, respectively. Due to using algorithmic thresholding, region localisation error can be slightly misleading between methods if a substantially different threshold is chosen as the level of threshold is generally correlated with increasing region localisation error. The approximate spacing between regions in the source space was around 10 mm, so an average region localisation error below 1 mm is insubstantial. Similar performance trends were observed across varying SNR levels and in the higher resolution source space of 20 484 vertices (see Fig S5), where geometric eigenmodes again exhibited a small advantage in cosine similarity and slightly better performance in explained variance and mean squared error.

##### Effect of using subject-specific vs. group-averaged structural data

For geometric eigenmodes, using subject-specific eigenmodes did not offer statistically significant improvement in RLE, RLE at 50% power or cosine score. For mean squared error and variance explained, there was a small improvement of *<* 1% that was statistically significant.

Using the group forward model resulted in a performance decrease of around 30% under perfect noise conditions but, surprisingly, a 4% increase at a SNR of 10 dB. It is possible that the smoothing of the brain, associated with group-averaged structural data, may have improved noise handling. For the connectome eigenmodes, using subject-specific eigenmodes reduced performance slightly (a cosine score of 0.74*±* 0.05 vs. 0.75 *±* 0.05), while using the group forward model dropped the cosine score by 40% to 0.44 *±* 0.03. Geometric eigenmodes appeared more robust to perturbations in subjects from the group-average.

##### Surrogate modes test

Both geometric and connectome eigenmodes offered statistically significant improvements over their surrogates for all metrics and for high SNR situations. For geometric eigenmodes, the cosine score was 0.76 *±* 0.07 compared to 0.28 *±* 0.07 for rolled surrogates and 0.75 *±* 0.07 for rotated surrogates. For connectome eigenmodes, the cosine score was 0.74 *±* 0.05 compared to 0.26 *±* 0.08 for rolled surrogates and 0.62 *±* 0.04 for rotated surrogates. The reason why geometric eigenmodes had a much smaller difference in performance compared to its rotational surrogates is due to the methodology of the rotational surrogates. Since the eigenmodes are rotated within their eigengroups (only applicable to geometric eigenmodes), the rotational surrogate method is significantly harsher for the geometric eigenmodes than the connectome eigenmodes, where it is equivalent to a form of spin test.

Additionally, applying the same surrogate test to the geometric eigenmodes constructed in a higher-dimensional source space (ico-5 spacing with 20 484 sources) revealed a more substantial difference in performance between the geometric eigenmodes and its rotational surrogates (see Fig S5). This suggests that the reduction in source space removed much of the finer-resolution spatial information that the geometric eigenmodes provided. Overall, the surrogate eigenmode tests suggest that there is physical meaning in the structural eigenmodes used and that both types were helpful in constraining the problem of source localisation.

### 3.3 Structural Eigenmodes on Localising the Epileptogenic Zone in Real Patients

As per Sun et al. [15], we localised the averaged interictal spike data of 20 patients who suffered from drug-resistant focal epilepsy and underwent resective surgery, where the resected region is treated as the base truth and compared to the estimate produced by the source localisation approaches. This is an imperfect source of truth, as areas that were active during interictal activity may lie outside of the resected region, and the resected region will be likely larger than the active area. Nevertheless, it is still suitable as a means to compare different source localisation approaches, as the resected region is highly likely to be the majority of the source of the interictal activity.

The geometric eigenmode approach had significantly higher recall compared to other methods at no expense to other metrics, region localisation error, recall, and ROCAUC (Receiver Operating Characteristic Area Under the Curve). Using the paired one-sided Wilcoxon signed rank test, comparisons had p-values of 0.0133 against connectome eigenmodes, 0.0068 against sLORETA and 0.0088 against eLORETA. This 30% improvement in recall against traditional approaches suggests that geometric eigenmodes have superior spatial coverage of the true source without overextending beyond the predominant region of interictal activity. An improvement in recall without penalty to other aspects in performance is an advancement compared to previous approaches, which needed to balance a trade-off between precision and recall between algorithms [15, 20, 59]. The relative performance increase of geometric eigenmodes against connectome eigenmodes mirror simulation results to suggest that group-level geometric eigenmodes should be preferred over using group-averaged connectomes.

## 4 Discussion

The presented results demonstrated that structural eigenmodes offer a computationally efficient means to estimate biologically feasible sources, with potential application to localising epileptic activity. Structural eigenmodes were able to represent simulated ictal activity at both source and sensor levels. Surprisingly, few modes were required to explain most of the variance in both source and sensor space, reflecting the fact that low wavelength modes dominate cortical activity[9, 60]. Both geometric and connectome eigenmodes demonstrated potential as efficient constraints for EEG/MEG source localisation, completing reconstructions within a fraction of the signal time. Importantly, both approaches provided similarly strong performance and were able to track the spread of a seizure, compared to mainstream approaches which were unable to do so. This is especially noteworthy for geometric eigenmodes given that the underlying simulations used the same subject-specific connectome as the connectome used to generate the connectome eigenmodes. Contrary to expectations that only a few modes would be required, the current study showed more modes significantly improved performance, most likely related to the ‘double descent’ phenomenon in machine learning literature [57, 58]. Finally, the geometric eigenmode approach was found to localise the epileptogenic zone successfully in epilepsy patients who had positive surgical outcomes, demonstrating significant alignment between the localisation results and the clinical findings.

### On the neurological meaning of eigenmodes

Previous work on fMRI has suggested that, relative to the entire brain space, both geometric eigenmodes and connectome eigenmodes offer a lower-dimensional and more interpretable view of the mesoscopic scale activity of the brain [9, 43]. This work focused on EEG, which provides a more direct measurement of neural activity and operates on temporal scales that closely align with neural dynamics, in contrast to the slower, indirect BOLD (Blood Oxygen Level Dependent) response measured by fMRI [61]. In simulated and real-world settings, both types of structural eigenmodes were able to find the most relevant spatial modes that reconstructed brain activity. This result held true under surrogate tests, in which the eigenmodes were altered in ways that preserved orthogonality or spatial structure. Hence, this work supports the view that both types of structural eigenmode offer neurologically meaningful interpretations of neural activity beyond acting as simple mathematical constructs.

Subject-specific modes offered performance similar to that of group-averaged modes in this analysis, suggesting that subject-specific modes were not vital to improving performance at the current resolution of 1000 cortical dipoles. This mirrors findings in the original paper, where, in high resolution fMRI data (32 492 vertices per hemisphere), individual-specific eigenmodes offered slight improvements compared to group-template derived eigenmodes in some individuals in reconstructing task-activation maps but not in reconstructing resting-state activity [9]. However, it is non-trivial to align subject and group-averaged spacing correctly. So, from a practical standpoint and to avoid alignment issues, algorithms should be run fully in subject space or group-averaged space. More importantly, weighting the eigenmodes based on wavelength was necessary for good source location performance, supporting previous findings that longer wavelength activity dominates cortical activity [9]. Future work can consider optimising the specific weightings, but the weighting schema chosen in this work based on physical principles performed very well.

### On the difference between geometric eigenmodes and connectome eigenmodes

When used as basis sets to constrain the source localisation problem of EEG, both geometric and connectome eigenmodes performed similarly across most metrics, with more noticeable differences emerging in mean squared error and variance explained. Given that the eigenmode weighting process, critical to the optimisation, was subtly different between the two approaches, and that the construction of the connectome and its eigenmodes require many procedural decisions, we do not consider the observed performance differences to be sufficient to argue for the superiority of one method over the other. Considering these results, we conclude that both geometric and connectome eigenmodes are substitutes in terms of performance, at least for the purposes of EEG/MEG source reconstruction. However, from a practical perspective, geometric eigenmodes only require the subject’s MRI scan, which is already required by the forward model, whereas connectome eigenmodes require additional data in the form of diffusion MRI and involve additional processing, which may affect the reconstruction results [37]. Indeed, there are complexities in aligning the source space with the streamline to parcellation mapping for connectomes, whereas aligning the source space in the geometric eigenmode approach is trivial. Furthermore, in the case where the subject-specific structural data is not available, our results suggest that geometric eigenmodes offer superior performance when using group-averaged data. As such, geometric eigenmodes may be preferred for the purposes of source localisation from a practical and ease-of-use standpoint.

### On biologically-informed constraints in source localisation

Traditional approaches have focused on mathematical arguments of smoothness and focality, and were less motivated by neurophysiology [8]. The issue of insufficient biological realism in source localisation problems is gaining attention in the field. Complementary studies have typically tried regularisation and other methods of incorporating connectome connectivity [62, 63] and cortical geometry [64] with some success. The other approach is best represented by the seminal work of Sun et al. [15, 28], who introduced the idea of biologically constraining neural networks with simulated data generated from dynamical neural models with great success. However, with deep learning approaches, a problem arises in validation because the simulated training and test data would be very similar, leading to the risk of over-fitting and low generalisation. This work shares the advantage of demonstrating complex source reconstruction using similarly complex models but ensures the source of the biological information is separate from the simulated data, making it more likely that this approach is generalisable to other settings. Deep learning methods continue to be an active area for source localisation research despite having the notorious requirement of needing to retrain the model on any kind of experimental change such as the electrode montage used [20, 21, 22]. In all, this study supports the view that the problem of source localisation is significantly aided by biologically informed constraints, and that both geometric and connectome eigenmodes provide relatively high gains for their mathematical simplicity and computational efficiency.

### On alternate formulation - regularisation

This work took advantage of the singular value decomposition of the Moore-Penrose inverse to arrive at the inverse solution. A regularisation approach to incorporate structural eigenmodes [63, 64] is also possible, which would align well with current Minimum-Norm approaches from the LORETA family. In such a scenario, the source space solution would instead be given by

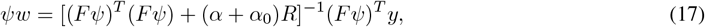

where *α* is the noise-dependent regularisation parameter, *α*_0_ is a regularisation constant to ensure numerical stability for all signal-to-noise ratios, and *R* is the regularisation matrix of choice. Examples of *R* include *I*, the identity matrix, for Tikhonov regularisation to arrive at a derivative of MNE, and *L*^*T*^ *L* where the Laplacian is used to arrive at a derivative of LORETA. Applying an *R* matrix dependent on *λ* to bias the eigenmodes, similar to the *W* weighting matrix in Eq. 8, would be appropriate for the eigenmode source localisation approach, where the power (*β*) of the eigenvalue can be tuned. The more general form of localisation using basis field expansions has been previously discussed [8, 19], with approaches using spherical harmonics [19] and, more recently, spherical head harmonics [18]. For our test cases, we found that the Moore-Penrose inverse was easier to work with, while suitable for a wide range of signal-to-noise ratios.

### On over-parameterisation in optimisation

The phenomena of overparmeterisation (also referred to as ‘double descent’ within the machine learning community) refers to the situation where a model has more parameters than necessary to fit the data, typically significantly more [58, 65]. Traditionally, this was expected to lead to overfitting, wherein the model would be dedicating parameters to fitting to noise instead of the true signal, leading to poor generalisability. However, modern deep learning challenges this view in that highly overparameterised models, namely deep neural networks, typically outperform everything else [57].

Within the context of eigenmode source localisation, Fig 5 showed that initially, as the number of parameters (modes) increased, performance increased as the model (algorithm) was able to output increasingly complex signals to express the true signal. Then, performance decreased with increasing number of parameters (modes) until the model (optimisation) crossed from the under-parameterised (*M < K*) to over-parameterised (*K < M*) regime (where we had more modes than sensors). From that point onwards, the metrics steadily increased. The poor performance when *M≈ K* was due to the system forcing a solution that fits the sensor signal exactly, but the EEG modes are increasingly aligned in EEG space, as seen in Fig 3b. When *M > K*, the increased solution space meant that there were infinite solutions and hence the minimum norm constraint was applied to return a unique solution. This is thought to have a regularisation effect to improve the reconstruction as the number of modes increased [65].

Future work should attempt to take advantage of the benefits and be mindful of the issues related to ‘double descent’. For example, incorporating loose orientations would increase the number of parameters by a multiple of 3, which may also lead to improvements explainable by overparameterisation, and the number of eigenmodes used should be far away from the number of independent sensors used.

### On the implications of tracking seizure propagation and localising the epilpetogenic zone in patients

From a clinical perspective, the ability to capture the dynamic evolution of seizure activity provides valuable insights beyond static localisation of the seizure onset zone. The spatial trajectory and timing of seizure spread may help disambiguate whether a region is truly epileptogenic and part of the origin or is instead part of the downstream propagation path. This distinction is critical for improving surgical planning because removing non-epileptogenic yet early-recruited regions may lead to unnecessary functional deficits without improved seizure control [66]. Additionally, being able to noninvasively model and reconstruct the spatiotemporal structure of seizures opens the door to more responsive and personalised interventions [49, 50]. For instance, real-time tracking of seizure spread could inform closed-loop stimulation paradigms that aim to disrupt propagation before clinical symptoms emerge [67]. In this context, the superior performance of structural eigenmodes in identifying recruited regions implies a potential role for these models in both pre-surgical evaluation and dynamic seizure tracking. With further improvements in source location and experimental technique, these models may even be able to test competing hypotheses of seizure generation and propagation – whether driven by fixed focal sources [68] or recruitment through travelling wavefronts [69, 70, 71].

In terms of better patient outcomes and quality of life, enhanced spatial resolution and interpretability may aid in distinguishing different seizure types, such as focal onset with rapid propagation vs. multifocal/generalised epilepsy, thereby refining seizure classification and informing more tailored treatment plans. More accurate localisation can reduce the need for extensive invasive monitoring, shorten time to surgery, and increase the likelihood of seizure freedom following resection [3]. At the same time, refined seizure classification would enable a more rational decision between pharmacological vs. surgical management strategies.

### On clinical integration and feasibility

Although the performance demonstrated by geometric eigenmodes is promising, successful clinical translation depends not only on accuracy, but also on interpretability and workflow compatibility. The output of the method, estimated source activation maps, can be visualised using standard cortical surface projections, similar to those already used in clinical neuroimaging such as CURRY. For localisation of epileptogenic zones within a cohort of people with epilepsy, geometric eigenmodes offered superior recall compared to widely used inverse solutions, with no compromise in precision or other performance metrics [16, 59]. This suggests that geometric eigenmodes could supplement, rather than replace, current noninvasive imaging techniques like PET or SPECT by refining hypotheses about seizure onset zones before invasive studies. Compared to approaches requiring individual diffusion MRI and tractography for each patient, geometric eigenmodes offer a lower-burden alternative. The finding that subject-specific eigenmodes were not essential implies feasibility in a wider variety of clinical situations, though having individual T1 MRI remains preferable for a more accurate forward model. As presurgical evaluation becomes increasingly improved by computational models and non-invasive imaging [31, 72, 73], tools like the geometric eigenmode approach may serve as a valuable addition to the clinician’s toolbox, bridging the gaps between observed clinical manifestations, patient brain structure, hidden functional seizure dynamics, and accurate patient prognosis. Future clinical validation would involve larger prospective cohorts, benchmarking against gold-standard outcomes such as Engel classification post-surgery, assessing real-time feasibility, and exploring integration with iEEG and surgical navigation platforms.

### On limitations

In this work, several algorithm parameters were fixed as part of the experimental design and assumptions. The depth weighting, which adjusts the forward model, was not applied. Dipoles were assumed to be fixed and oriented perpendicular to the cortical surface, but the orientation could have been loose. Care must be taken when considering loose orientations, as results can be misleading as discussed in Sec.S1.5. The choice of the *β* parameter for eigenmode weighting as a function of the eigenvalues (Eq. 8) was based on physical arguments, rather than optimisation. Similarly, the steps involved in the construction of the connectome, such as spatial smoothing and thresholding, relied on expert judgement. Although all of these parameters can be adjusted and tuned in practice, extensive tuning of hyperparameters risks overfitting to a specific dataset and may compromise the generalisability of the methodology.

Another main limitation was the reliance on simulation as the source of truth for evaluation. While simulations provide valuable ground truth for assessing methodological performance, they inevitably simplify the complexity of real neural dynamics and are unlikely to capture the full range of noise and variability present in real-world data. Specifically, while the Epileptor model is considered state of the art and has a strong track record of reproducing many spatial and temporal features of epileptic dynamics at both microscale and macroscale levels [30, 48, 50], its canonical form captures only one of the twelve theoretically defined dynamical transitions associated with seizure activity [74]. Moreover, assumptions made during simulation design, such as source extent, timescales, and propagation dynamics, may introduce biases in favour of specific modelling approaches. For example, the reliance of the coupled neural mass model on the structural connectome may advantage the connectome eigenmode approach by construction. Nevertheless, while absolute accuracy in simulation often overestimates performance in real data, relative performance trends, such as the comparative ranking of methods or the impact of specific parameter adjustments, often generalise.

To address these concerns, we complemented the simulation-based evaluation with application to real patient data from individuals who underwent surgical resection. However, it’s important to reiterate that from a clinical perspective, the localisation of interictal spikes does not necessarily reflect the true epileptogenic zone, which may confound the evaluation and risks over-interpretation. This highlights the need for future work to systematically validate the approach in prospective studies across diverse patient populations, ideally incorporating multi-modal data such as fMRI or intracranial recordings to better assess clinical utility and confirm generalisability. Although simulations cannot substitute for clinical validation, in domains like source localisation where a definitive ground truth is currently not available, they serve as a critical intermediary, helping to guide expectations, identify limitations, and narrow the space of viable methods prior to clinical deployment.

## 5 Conclusion

Source localisation is an important technique for improving the spatial accuracy of EEG and MEG measurements [5]. Improved algorithms will lead to an improved understanding of the brain, such as advances in our knowledge of consciousness [75], and has direct applications in the diagnosis and treatment of neurological disorders. This work combined the recent pivotal idea in computational neuroscience of structural eigenmodes to biologically constrain the optimisation problem of source localisation and found it to be highly effective in representing neurologically plausible signals in terms of its accuracy, computational efficiency, and ease of use. Furthermore, the presented approach showed promise in noninvasively analysing the origins of ictal and interictal activities, which may one day drive improved outcomes for people with epilepsy.

## Code Availability

All code for the source localisation pipeline can be found at: https://github.com/Spokhim/sesl. All code should be easily compatible with current MNE-Python pipelines whether it is EEG or MEG. If there is enough interest, the author may look into a proper implementation into MNE-Python.

## Acknowledgements

The author would like to thank the Australian neuroimaging and computational neuroscience community for their encouragement and support, as well as the international friends met during travel in the Americas. This research was supported by The University of Melbourne’s Research Computing Services, the Petascale Campus Initiative, and an Australian Government Research Training Program (RTP) Scholarship. This research was funded by the Australian Research Council (Discovery Project DP200102600).

## Notes

### Competing Interest Statement

The authors have declared no competing interest.

