## Supplementary Document for "Structural Eigenmodes of the Brain to Improve the Source Localisation of EEG: Application to Epileptiform Activity"

### S1 Supplementary Material

#### S1.1 Simple Source Simulations

For a more direct comparison with previous literature which focuses typically on one to few sources of various extent [17, 18, 19, 20, 21, 22], we also had a simple set of single-source simulations. These sources were generated with varying sizes such that for each simulation, one cortical region of the 1000-region parcellation was selected as the centre. This simulation would also be more similar to the case of averaged, interictal spikes. We compared the performance of geometric eigenmodes without weighting, with weighting, and against eLORETA. The geometric eigenmode approach had superior performance to eLORETA at the single-patch source level. The significant improvement in metrics with weighting mirror previous results stating that low-wavelength activity dominate cortical activity [9].

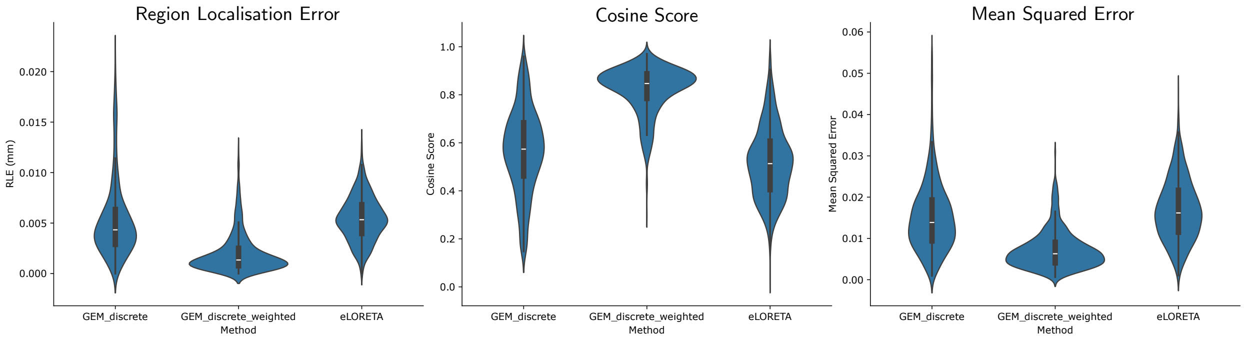


Figure S1: Patch simulations demonstrating performance of: i) GEMs with no weighting, ii) GEMS with eigenvalue weighting, iii) eLORETA. Sources had fixed orientations.

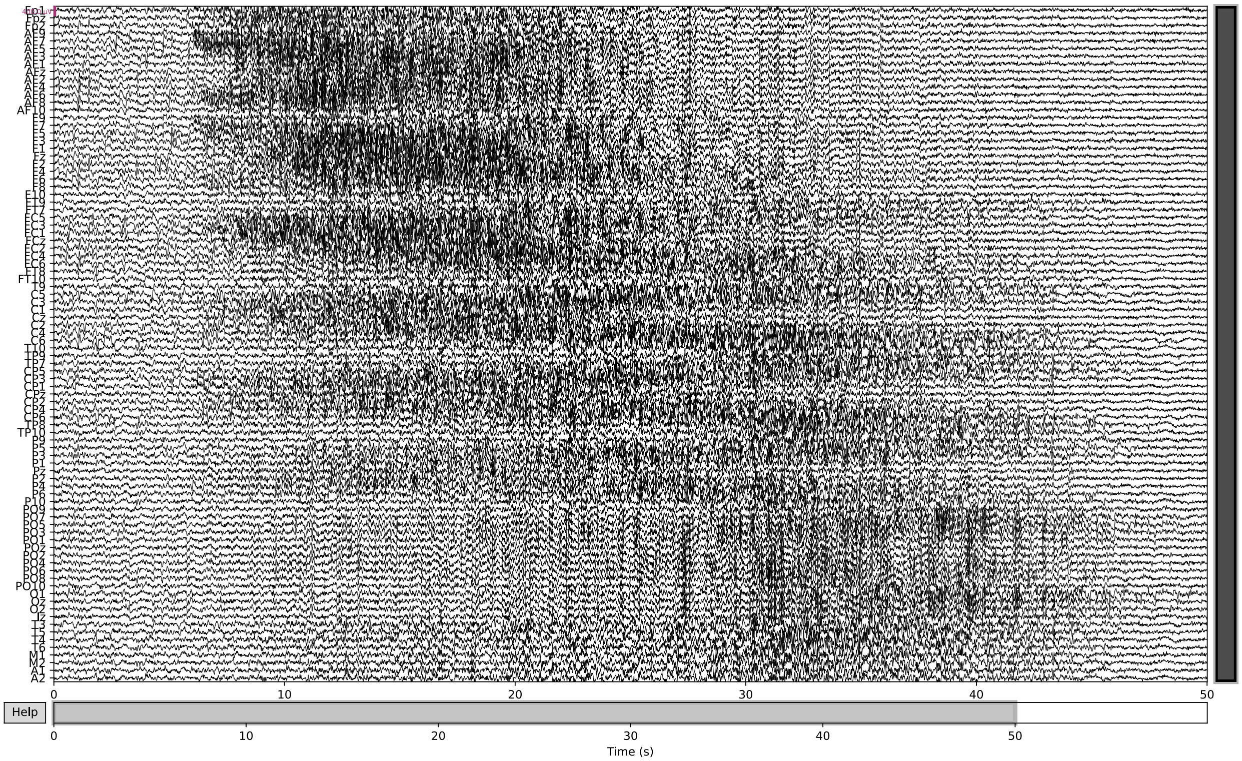


Figure S2: EEG trace of simulation with SNR of 10 dB. The simulated EEG exhibited clinically consistent features, including focal onset, spatial spread, and temporal evolution with characteristic slowing.

##### S1.2 Connectome Eigenmode Weighting and Combined Structural Eigenmodes

Per Eq. 8, geometric eigenmodes were weighted by $\beta = 1/2$ and connectome eigenmodes were weighted by $\beta = 1$ so that the weights ended up being very similar after normalisation as demonstrated in Fig.S3. The regularisation constant was selected to be the same for both to ensure that the overall weights were similar. These are the weights used when testing geometric and connectome eigenmodes together as the basis set for source localisation, which does not appear to offer a substantial difference (see Fig.S4). Explicitly, Fig S3 is graphing:

$$W_G = \frac{\pi}{0.1 + |\frac{\lambda_G}{\bar{\lambda}_G}|^{1/2}} \tag{18}$$

$$W_C = \frac{\pi}{0.1 + |\frac{\lambda_C}{\bar{\lambda}_C}|} \tag{19}$$

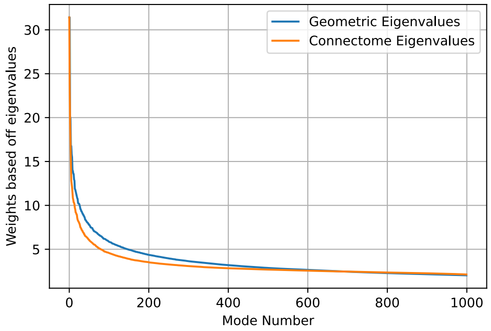


Figure S3: Visualising the weights after eigenvalue weighting. Noting that a different power for the eigenvalue was applied so that the GEMs and CEMs have similar weights.

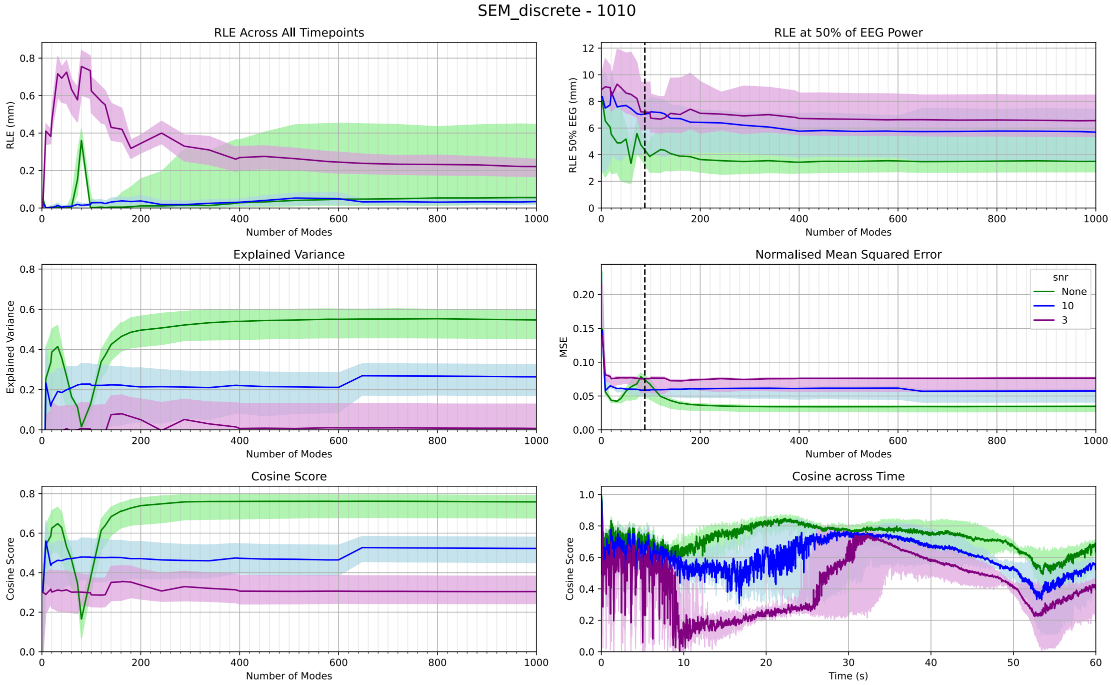


Figure S4: Performance for the structural eigenmode approach when concatenating the set of modes of geometric and connectome eigenmodes after weighting as per Fig.S3.

##### S1.3 High-density Source Space

Within the main text, the analysis focused on a 1000-vertices source space which matched the spatial resolution of the underlying discrete connectome. Here we present analysis of the structural eigenmode source localisation approach performed on icoshedron-5 (ico-5) source spacing consisting of 20 484 dipole sources. In ico-5 space, multiple dipoles are associated with each cortical region and the underlying source time series are projected from 1000 cortical regions to the 20 484 sources. Furthermore, Laplacian smoothing of neighbouring values is applied with 5 iterations to remove boundary effects of the projection. The corresponding EEG signal is then calculated with a forward transform from the ico-5 space. To perform high-resolution connectome eigenmode analysis, the eigenmodes were constructed on a high resolution connectome in fsLR space and subsequently mapped to ico-5 source spacing [36, 76].

Both the geometric eignemodes and the connectome eigenmodes performed reasonably well in this high-density source space, with similar trends found in the 1000-vertices source space. At this high-density source space, a more notable distinction between geometric eigenmodes and its rotational surrogates emerges, and the connectome eigenmodes perform better than the geometric eigenmodes' rotational surrogates. See Fig. S5 for the comparison of metrics, and Fig. S6 for a visualisation of the source estimates.

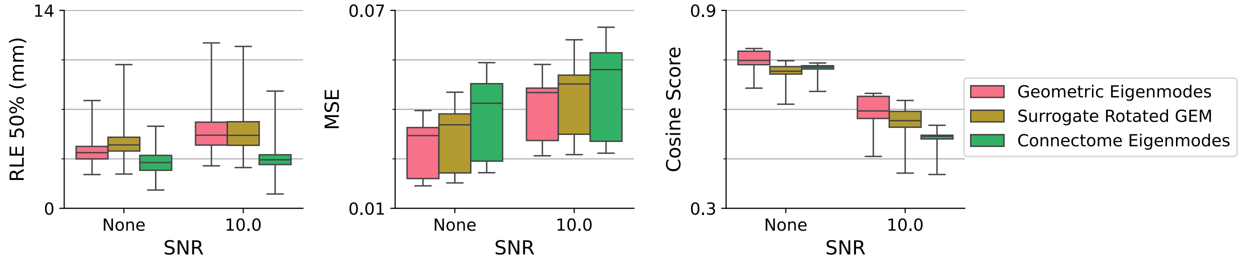


Figure S5: Structural eigenmode approach performed on ico-5 source space consisting of 20 484 sources. In ico-5 space, multiple dipoles are associated with each cortical region and the underlying source time series are projected from 1000 cortical regions to the 20,484 sources. The corresponding EEG signal is then calculated with a forward transform from the ico-5 space. Both GEMs and CEMs performed reasonably well in this high density source space.

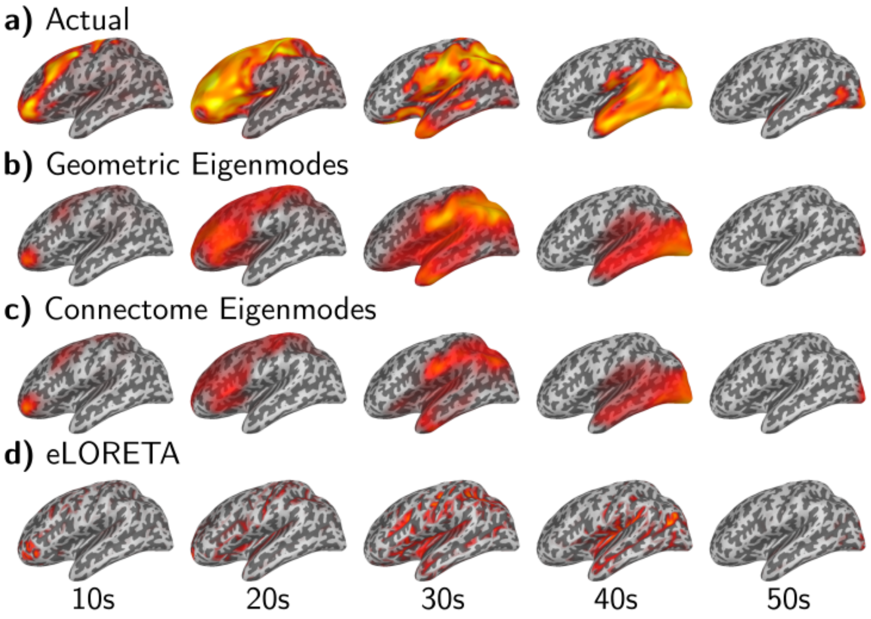


Figure S6: Source localisation example for seizure originating from thalamus plotted on left hemisphere in ico-5 source space consisting of 20 484 sources.

##### S1.4 Different Number of EEG Electrodes

Within the main text, we focused our analysis on 10-10 spacing corresponding to 88 EEG electrodes as this is a common number used in research. We have also investigated 10-05 spacing corresponding to 343 electrodes (Fig. S7 and 10-20 spacing corresponding to 21 electrodes (Fig. S8).

Key highlights would be both geometric eigenmodes and connectome eigenmodes were able to reasonably resolve sources at all electrode configurations. The algorithms did not perform well at a SNR of 3. At 21 and 88 electrodes, the MN methods were unable to accurately determine the polarity of sources, as evidenced by the variance explained being close to 0, and the relatively low cosine score. At 343 electrodes, the MN methods were able to resolve the polarity of sources, leading to substantial increases in metrics.

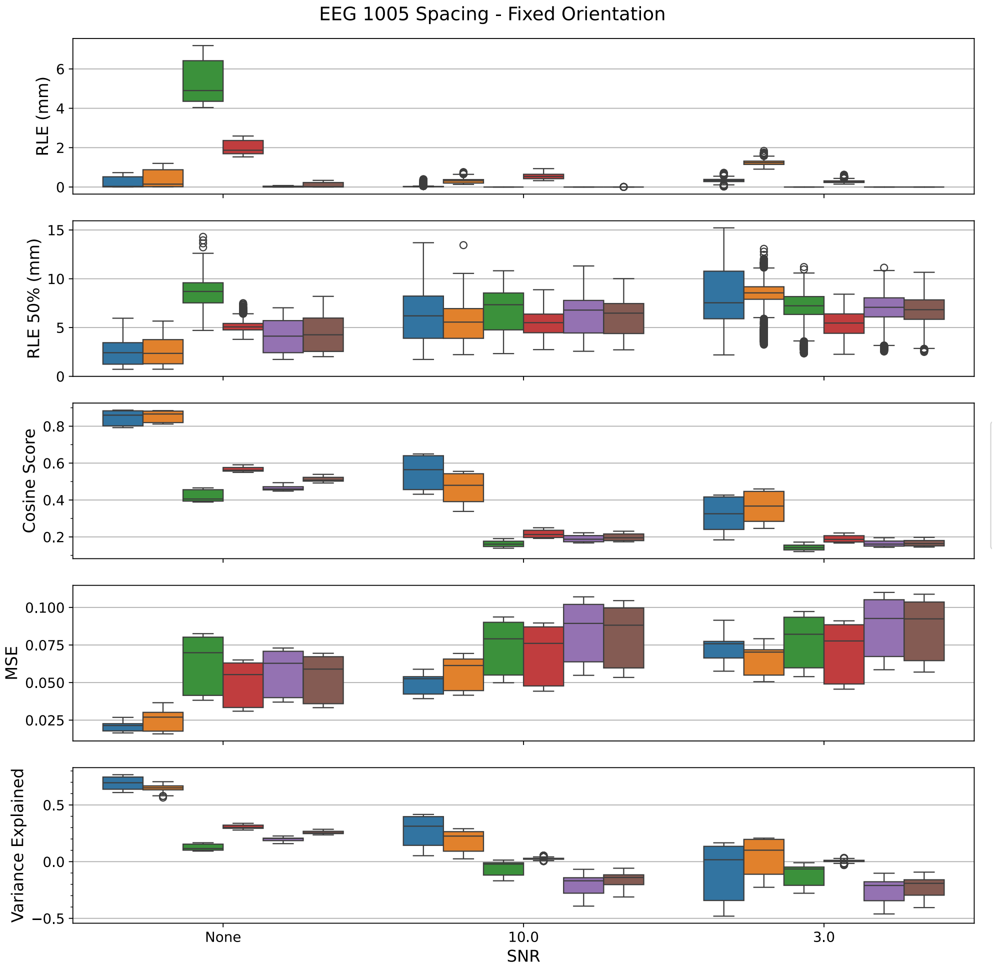


Figure S7: EEG 10-05 spacing metrics for fixed orientation. Legend: Blue = GEM, Orange = CEM, Green = dSPM, Red = MNE, Violet = sLORETA, Brown = eLORETA.

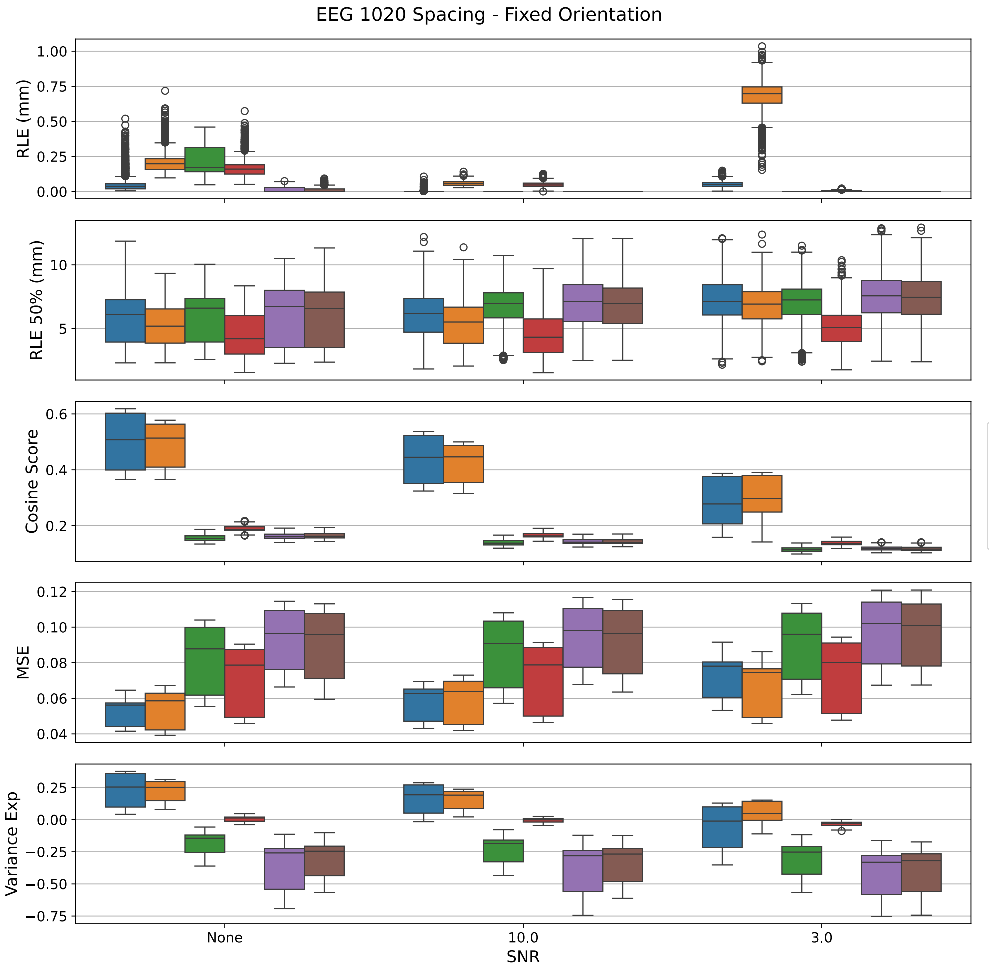


Figure S8: EEG 10-20 spacing metrics for fixed orientation.

##### S1.5 Allowing Loose Orientations of Dipoles

Allowing the orientation of the dipole to be free to deviate from the normal of the cortical surface (also referred to as loose orientation) is a common approach in source localisation [46, 77]. As previously stated, in this work the dipoles were oriented normal to the cortical surface for the simulation. However, preliminary results suggest that allowing a deviation from the normal of the cortical surface, by setting the 'loose' parameter to $! = 0$, increased performance for all source localisation methods, and the LORETA family more significantly. The standard explanation would be that the loose orientation provides extra degrees of freedom to account for small errors with the forward model, mesh discretisation, downsampling, etc. This section is to provide evidence that such a performance increase can be deceptive and that the performance increase is primarily a result of condensing the vector source estimate into a magnitude measurement, rather than any error correction.

First, let us demonstrate how setting a loose orientation appears to improve source localisation performance. This is as seen by comparing Fig.6 with Fig.S9. For the EM methods, this improved the handling of noise, but decreased performance slightly under no noise conditions. For the LORETA family, this significantly improved overall performance for all SNRs.

However, the apparent improvement in performance can be demonstrated to be an artifact of vector estimation. After estimating the loose source estimate, there is a need to compare it with the true source estimate which is oriented normal to the cortical surface. The typical method, as default in MNE-Python, is to condense the vector into a magnitude and compare, and this is what was performed to generate Fig. S10c)-d) and Fig. S11c)-d). Both figures plot source estimates from a single simulation (seizure originating from Thalamus at $g = 2$). Sequentially, a) represents fixed

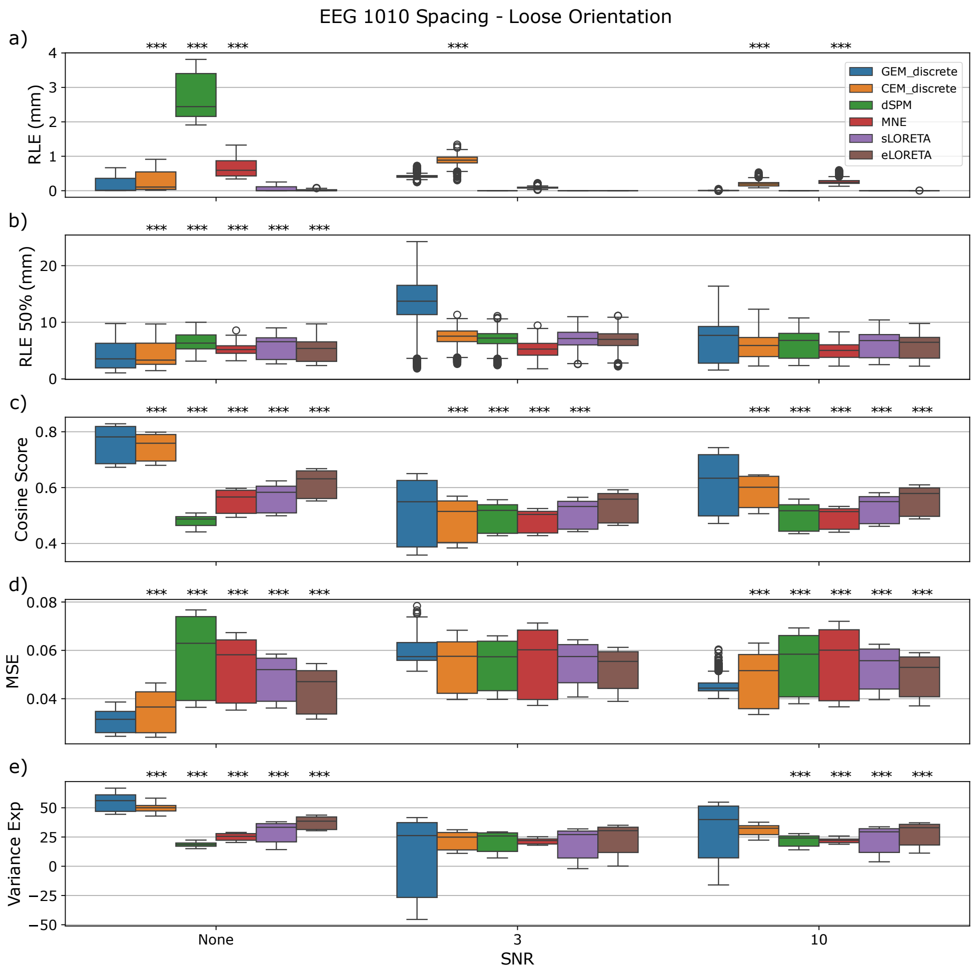


Figure S9: Metrics for loose orientation

orientation, b) represents fixed orientation but then taking the absolute value of source estimate, c) represents setting a loose orientation of 0.05, and d) represents loose orientation of 1. It can be seen that under these conditions, adding the loose parameter significantly increased performance. Further increase of the loose parameter appears to slightly increase the performance for eLORETA. Fig.S11b) demonstrates that the performance increase can also be seen by simply taking the absolute value of the fixed orientation estimate. Essentially, what this means is that the removal of the direction (and only taking the magnitude) is producing the bulk of the performance increase, rather than the allowance of deviation to the cortical surface. Indeed, Fig.S11e) displays the estimate and metric when only considering the orientation component normal to the surface results in relatively poor performance. This is why in the main text we present results in the fixed orientation source space.

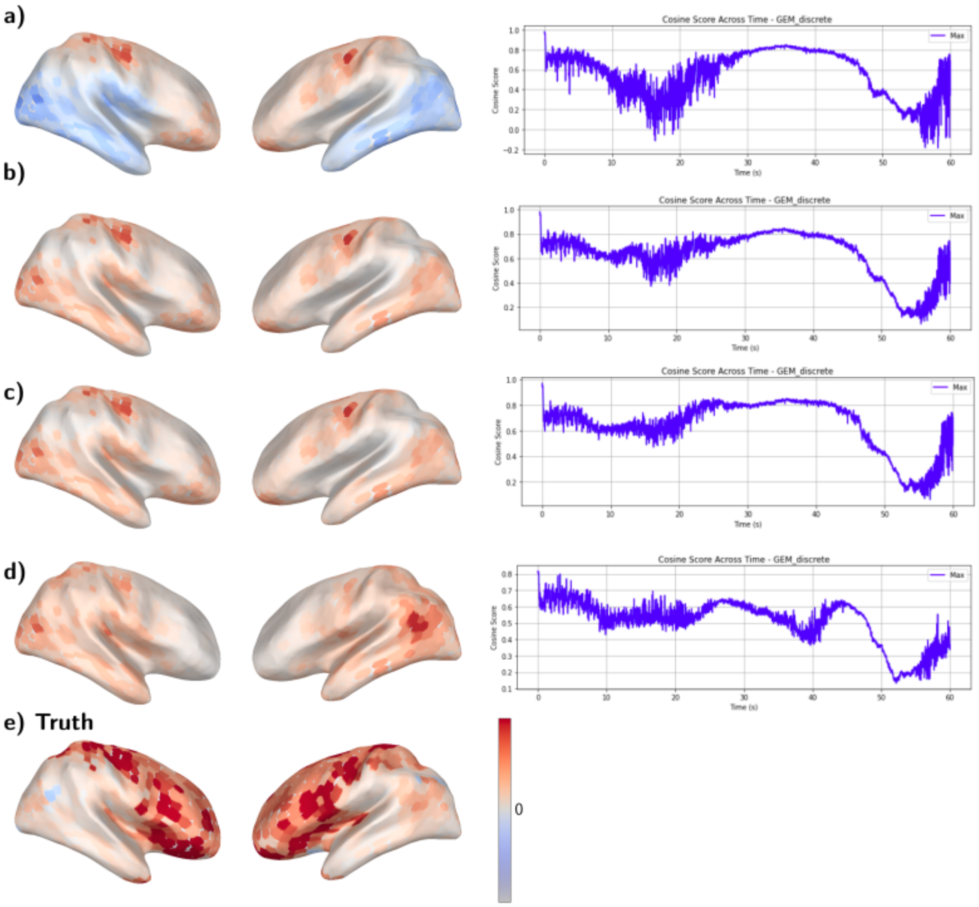


Figure S10: GEM method adjusting the loose parameter of a thalamic seizure with SNR 10. The localisation result at t=20s is visualised on the brain surface as it is a timepoint of unstable and relatively poor performance. The cosine score across the entire simulation is plotted beside. a) Loose = 0 (fixed orientation). b) Loose = 0, with np.abs(estimate). c) Loose = 0.05 (loose orientation). d) Loose = 1 (free orientation). e) True source, noting that the colourscale is diverging, whereas main text plots have been one-sided. Comparing a) with b), under noisy conditions, the algorithm may incorrectly estimate the correct polarity. Between 50s and 60s, a few sparse sources are active, and noise causes the algorithm to estimate the location incorrectly, noting that cosine score does not account for spatial similarity in that an estimate that is off by 1cm is treated the same as if it was off by 10cm.

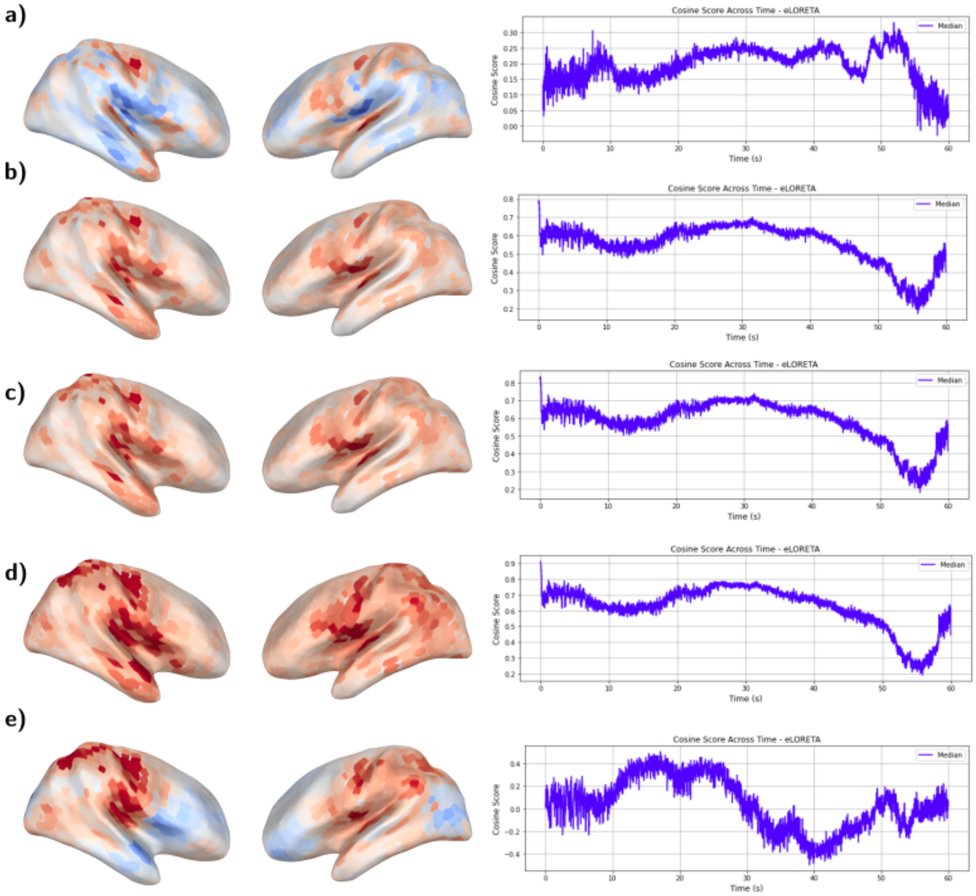


Figure S11: eLORETA method adjusting the loose parameter of a thalamic seizure with SNR 10. The localisation result at t=20s is visualised on the brain surface. The cosine score across the entire simulation is plotted beside. a) Loose = 0 (fixed orientation). b) Loose = 0, with np.abs(estimate). c) Loose = 0.05 (loose orientation). d) Loose = 1 (free orientation). e) Loose = 1 (free orientation), but taking only the normal to the surface component.

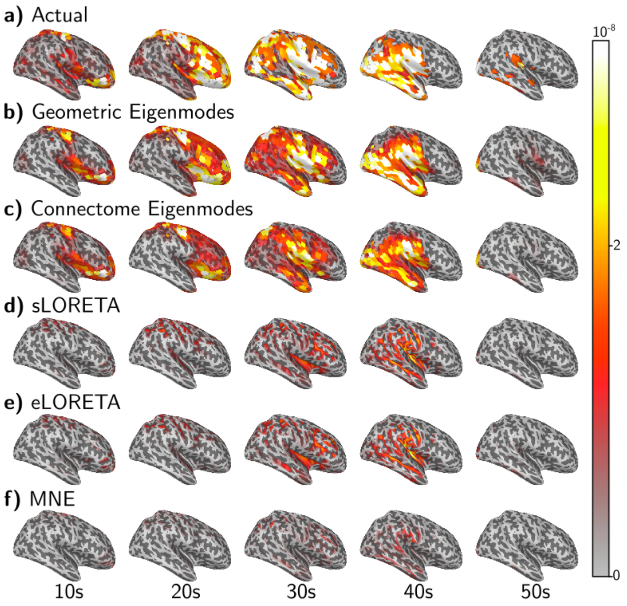


Figure S12: Source localisation example for seizure originating from thalamus plotted on right hemisphere 5 time points are selected in increments of 10s and visualised on the right hemisphere. 50% EEG power for this simulation occurs at the 30s. The colour scale is set such that maximum amplitude is white, progressing to yellow, orange, red, and transparent for 0. a) The actual or "ground truth" source activity as generated from the Epileptor model. We see that as time progresses, the seizure spreads from the anterior to posterior regions of the brain. During this time, different areas of the brain are activated in varying spatial extents and patterns. b) & c) Both geoemtric eigenmodes and connectome eigenmodes are able to perform reasonably well in sparse activations (10s and 50s), as well as dispersed activations (20s - 40s). d)-f) The LORETA family, predict more focal sources with default parameter settings and regularisation parameter set according to the $10^{-SNR/10}$ heuristic.

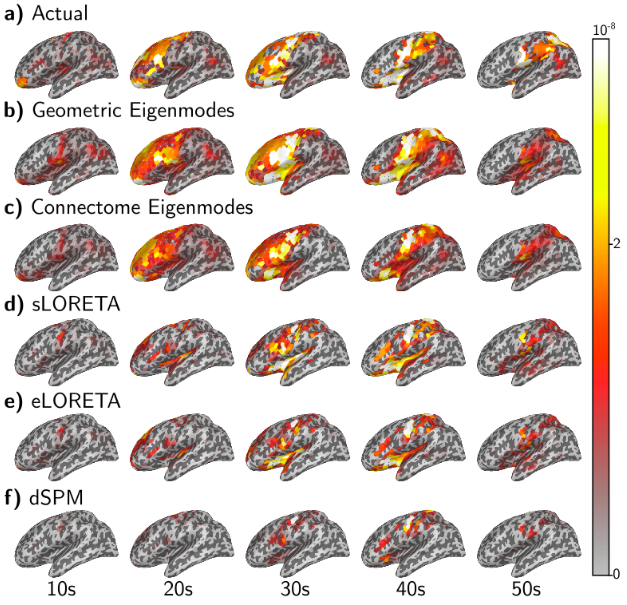


Figure S13: Seizure originating from cingulate cortex and partial spread ($g = 1.5$). All methods are kept at fixed orientation with no depth weighting.
